# Growth rate alterations of human colorectal cancer cells by 157 gut bacteria

**DOI:** 10.1101/2019.12.14.876367

**Authors:** Rahwa Taddese, Daniel R. Garza, Lilian N. Ruiter, Marien I. de Jonge, Clara Belzer, Steven Aalvink, Iris D. Nagtegaal, Bas E. Dutilh, Annemarie Boleij

## Abstract

Several bacteria in the human gut microbiome have been associated with colorectal cancer (CRC) by high-throughput screens. In some cases, molecular mechanisms have been elucidated that drive tumorigenesis, including bacterial membrane proteins or secreted molecules that interact with the human cancer cells. For most gut bacteria, however, it remains unknown if they enhance or inhibit cancer cell growth. Here, we screened bacteria-free supernatants (secretomes) and inactivated cells of over 150 cultured bacterial strains for their effect on CRC cell growth. We observed family-level and strain-level effects that often differed between bacterial cells and secretomes, suggesting that different molecular mechanisms are at play. Secretomes of *Bacteroidaceae*, *Enterobacteriaceae,* and *Erysipelotrichaceae* bacteria enhanced CRC cell growth, while most *Fusobacteriaceae* cells and secretomes inhibited growth, contrasting prior findings. In some bacteria, the presence of specific functional genes was associated with CRC cell growth rates, including the virulence genes TcdA in *Clostridiales* and FadA in *Fusobacteriaceae*, which both inhibited growth. *Bacteroidaceae* cells that enhanced growth were enriched for genes of the cobalamin synthesis pathway, while *Fusobacteriaceae* cells that inhibit growth were enriched for genes of the ethanolamine utilization pathway. Together, our results reveal how different gut bacteria have wide-ranging effects on cancer cells, contribute a better understanding of the effects of the gut microbiome on the human host, and provide a valuable resource for identifying candidate target genes for potential microbiome-based diagnostics and treatment strategies.

## INTRODUCTION

Over the past few decades, thousands of microbial species have been identified in the human gut microbiome. Each person harbors an estimated 3.8 x 10^13^ bacteria in their guts with a relatively stable species composition (1–3). Changes in the composition of the gut microbiome have been linked with various diseases including colorectal cancer (CRC), the third leading cancer worldwide with an incidence that is estimated to rise in the coming decennium (4, 5). Besides important environmental and genetic factors, the initiation and development of CRC is affected by the human gut microbiome. During tumorigenesis, the bacterial composition on the affected intestinal mucosa changes in a process that is described by the bacterial driver-passenger model (6, 7). Bacterial drivers may facilitate the acquisition of hallmarks of cancer (8) in mucosal cells by generating DNA-damage, reducing the epithelial barrier function, and stimulating pro-carcinogenic immune responses. As the tumor microenvironment changes in the transition from adenoma to carcinoma, bacterial passengers compete with the drivers, further shifting the microbiome towards a CRC signature (9, 10). The newly acquired bacteria may in turn affect tumor development, by potential inhibiting, enhancing, or neutral effects on tumor cell growth (6). These CRC-specific changes offer the opportunity to use the bacterial community for diagnostic, prognostic, preventive, and treatment measures (11–20).

In faeces of CRC patients, the genera *Fusobacterium*, *Peptostreptococcus, Ruminococcus*, and *Escherichia/Shigella* are enriched, while *Bacteroides* and *Lachnospiraceae* (*Roseburia, Lachnospira, Anaerostipes*) have a lower relative abundance (5, 9, 11, 21–25). Even though phyla such as *Bacteroidetes* are depleted, single species like enterotoxigenic *Bacteroides fragilis* may be enriched in some CRC patients (25–27). Comparing tumor tissue with adjacent normal tissue on the intestinal mucosa of CRC patients revealed that *Fusobacterium, Prevotella, Streptococcus*, and *Peptostreptococcus* were enriched in tumors while *Blautia, Escherichia, Pseudomonas*, and *Faecalibacterium* were enriched in the normal mucosa (5, 7, 9, 11, 22–24, 28).

Many different bacteria have been associated with CRC tumors (10), but we are only beginning to understand the different mechanisms that are involved. The effect of bacteria on CRC cell growth may be driven by direct bacterial-to-epithelial cell-cell contact or by secreted products present in the secretome (29, 30). Membrane-bound bacterial proteins that require cell-cell contact can activate epithelial cell signalling. For example, the passenger bacterium *Fusobacterium nucleatum* encodes the membrane protein *Fusobacterium* adhesin A (FadA) that binds to E-cadherin, activating β-catenin signaling and resulting into increased tumor growth (31–33). Specific *E. coli* species with *eae* gene express the adhesin protein intimin on their membrane surface which bind and cause lesions to gut epithelial cells, allowing bacteria to breach the colonic barrier. Once the bacteria are bound to the epithelial cells, this allows them to inhibit DNA repair proteins, further contributing to lasting DNA damage (34–36). Several secreted bacterial toxins are known that can bind to receptors or pass through the cell membrane into the cytoplasm. The driver bacterium enterotoxigenic *B. fragilis* secretes the metalloprotease *B. fragilis* toxin (BFT) which leads to E-cadherin cleavage and increased wnt-signalling in colon epithelial cells and to tumor formation in mice and increased cell proliferation *in vivo* (37, 38). Bacteria such as *Escherichia coli* containing the *pks* island are able to produce a genotoxin called colibactin. Upon mucosal breach, colibactin reaches the epithelial cells and alkylates DNA, ultimately leading to tumorigenesis (39–41). These examples show that taxonomically diverse bacteria may lead to the acquisition of hallmark capabilities via diverse mechanisms.

The aim of this study was to examine the effects of bacterial cells and their secretomes on the growth rates of epithelial cells. We tested the effect of 157 different gut bacteria on the growth rates of five CRC cell lines and one non-cancerous kidney cell line. Our results revealed that different bacterial families specifically inhibit or enhance CRC cell growth, although contrasting effects could be observed between some closely related strains. Both known virulence genes and novel microbial pathways were associated with the different growth rate changes. These results provide the first large-scale *in vitro* analysis of the effects of different microbial strains on epithelial cell growth.

## MATERIALS AND METHODS

### Bacterial strains

We selected 116 different gut bacteria, including species that are depleted or enriched in CRC patients whose genome sequences were available in the PATRIC database (42). Additionally, we selected specific bacteria without sequenced genomes (n=39) that were strongly linked to CRC or were isolated from CRC tissue, including *Streptococcus bovis* (43, 44), *Clostridium septicum* (44–46), *Clostridioides difficile* (47, 48)*, Bacteroides* sp. (27), and the two potentially beneficial bacteria *Lactobacillus casei Shirota* (49–51) and *Akkermansia muciniphila* ATCC BAA-835^TM^ (52–55). Together, we analysed 157 bacterial strains isolated from the human gut microbiome (Supplementary Table S1).

We purchased 96 bacterial strains from the reference catalogue of the Human Microbiome Project (HMP, Prof. Dr. Emma Allen-Vercoe from the University of Guelph in Guelph, Canada); 24 bacteria were kindly provided by Prof. Dr. Cynthia L. Sears from Johns Hopkins Medical Institutions, Baltimore, MD, USA; five strains were purchased from DSMZ (*Clostridium septicum* (Macé 1889) Ford 1927 DSM7534, *C. difficile* (Hall and O’Toole 1935) Lawson et al. 2016 DSM27543 (known as *Clostridium difficile* 630, PMID: 6870225), *Fusobacterium nucleatum* Knorr 1922 DSM15643, *Fusobacterium nucleatum* subsp. *polymorphum* (ex Knorr 1922) Dzink et al. 1990 DSM20482, and *Peptostreptococcus stomatis* Downes and Wade 2006 DSM17678); one strain from ATCC (*Streptococcus agalactiae* ATCC13813); and 31 bacteria were in stock at the Radboud University Medical Center in Nijmegen, The Netherlands (56–58). Information about the bacterial strains, their origin, growth media, and their genome sequence is provided in Supplementary Table S1.

Bacteria were cultured in their respective media under anaerobic conditions at 37°C aired with nitrogen gas (N_2_, see Supplementary Table S1). Alternatively, *Ralstonia* sp. 5_2_56FAA, *Ralstonia* sp. 5_7_47FAA and *Pseudomonas* sp. 2_1_26 were grown aerobically at 37°C, and *Bacillus smithii* was grown anaerobically at its preferred temperature of 50°C. Eight media were prepared to culture bacteria (Supplementary Table S1): brain heart infusion-supplemented (BHI-S, ATCC Medium 1293), BHI-S supplemented with 0.1% Tween 80 at pH 8.0 (BHI-T((59), NADC - 99X medium (ATCC Medium 1804), Desulfovibrio medium (ATCC Medium 2755), Modified Reinforced Clostridial Medium (RCM, ATCC Medium 2107), Reinforced Clostridial medium with sodium lactate (60% solution) at a concentration of 1.5% (RCM+, from ATCC Medium 1252), differential RCM (DRCM), and nutrient broth (NB, ATCC Medium 3). Bacteria were grown until the medium was turbid and the optical density (OD) was at least 1.0. Absorbance was obtained by measuring OD at 600nm.

### Obtaining secretomes and bacterial cells

To investigate the effect of secreted molecules, we obtained secretomes from 154 bacterial cultures as follows. After culturing, bacteria were centrifuged at 4,700 rpm for 10 minutes and some bacteria subsequently at a higher speed to settle them down (see Supplementary Table S1). Next, supernatants were spun at 4,700 rpm for 10 minutes to pellet any remaining bacteria and filter-sterilized using 0.2 µm filters (Sigma-Aldrich, USA). Molecules larger than 10 kDa were concentrated using amicon ultra-15 centrifugal filters (Merck Millipore, Merck, USA). Concentrated supernatants were frozen at −80°C until further use and are further referred to as secretomes.

To investigate the effect of bacterial surface-bound molecules, 145 bacteria were inactivated to allow cellular growth rates to be assessed by measuring metabolic activity (see the section “MTT assay” below). Cells of twelve spore-forming bacteria under our experimental conditions were excluded from this analysis (see Supplementary Table S1). Pelleted bacteria were resuspended in 70% ethanol and incubated for 5 minutes according to our inactivation protocol (60). After centrifugation at 20,000 rcf for 30 seconds, bacteria were washed in PBS twice and inactivated bacteria were frozen at −80°C until further use.

### Culturing of cell lines and identification of their mutational landscapes

Five CRC cell lines (Caco-2, HCT15, HCT116, HT29, and SW480) and one non-cancerous embryonic kidney cell line (HEK293T) were selected based on their differences in mutational landscapes (61). Known oncogenes and tumor suppressor genes were confirmed with targeted single molecule molecular inversion probe (smMIP) mutation analysis as described (62), with additional smMIPs targeting CXCR4 (NM_001008540.1) codons 281-357, EZH2 (NM_00004456.4) codons 471-502, 618-645, 679-704, and SF3B1 (NM_012433.2) codons 603-471, 833-906 (smMIP sequences are available upon request), at the Radboud Diagnostics Department, Nijmegen, The Netherlands (Supplementary Table S2). Cells were split twice a week using Dulbecco’s Modified Eagle Medium (DMEM, Lonza, Switzerland) supplemented with 10% fetal bovine serum (FBS, Thermo Fisher Scientific, USA) and 1% Penicillin/Streptomycin (Pen/Strep, Thermo Fisher Scientific, USA). Cells used for this study were not split for more than 25 passages.

For MTT assays, optimal cell seeding in 96 well plates (Greiner Bio-One, Austria) was defined at 5,000 cells/well for Caco-2, 4,000 cells/well for HCT15, 1,500 cells/well for HCT116, 10,000 cells/well for HT29, 6,000 cells/well for SW480, and 20,000 cells/well for HEK293T. Cells were incubated overnight to attach before addition of secretomes or bacterial cells.

### MTT assay

Secretomes were diluted to their original concentrations in DMEM supplemented with 10% FBS and 1% Pen/Strep. Inactivated bacteria were diluted with DMEM supplemented with 10% FBS and 1% Pen/Strep to an OD of 0.2 and incubated on cell lines up to 72 hours. For secretomes, filter sterilized bacterial culture media used to generate secretomes served as a control for cell growth in the assay, while for bacterial cells, DMEM supplemented with 10% FBS and 1% Pen/Strep served as control for cell growth. MTT assays were performed to measure the cell metabolic activity every 24 hours. These assays involve the conversion of the water-soluble yellow dye MTT [3-(4,5-dimethylthiazol-2-yl)-2,5-diphenyltetrazolium bromide] to the insoluble purple formazan by the action of mitochondrial reductase. At 24, 48 and 72 hours 10 µl of MTT (5 mg MTT dissolved per mL PBS) dye was added to the wells and cells were incubated for 3 hours. Formazan was then solubilized with 150 µL MTT solvent (0.5 mL of 10% Nonidet (dissolved in water), 50 mL isopropanol (2-propanol) and 16,7 µL hydrochloric acid fuming 37% together) and the concentration determined by optical density at 570 nm (BioRad microplate reader model 680). Metabolic activity of cell lines was used as a measure for cell growth (63). All experiments were performed in quadruplicate.

#### Cell growth analysis

Absorbances at optical density of 570 nm were measured by MTT assays at four time points (0, 24, 48, and 72h). Growth rates per hour were calculated by dividing the difference in absorbance by the time interval between measurements (24 h). Results from four independent experimental replicates were averaged. Growth rates between 24-48 h were selected for comparative analysis because that time window captured the optimal growth for most cell lines (Supplementary Figure S1).

A growth rate score was defined by subtracting the growth of negative controls (cells cultured without bacterial products) from those cultured with bacterial products (secretomes or bacterial cells). Scores greater than 0 refer to enhanced growth, while scores lower than 0 refer to inhibited growth (see Supplementary Tables S3 and S4). To allow comparability between cell lines, normalized growth rate scores were calculated by dividing per cell line the scores (g) of each of the n bacteria by the norm (|g|) of the experimental condition 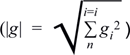. The normalized growth rate scores were used to evaluate whether the enhancing or inhibiting effect of a bacteria is significantly stronger than the mean effect observed for all other bacteria. For this purpose, growth rate scores were transformed into z-scores and p-values were computed by separate one tailed z-tests for enhancing or inhibiting effects. Based on these p-values, strains were identified as significant enhancers or inhibitors of growth by using a significance cut-off of p<0.05.

The overall effects of bacterial secretomes and inactivated cells were visualized by projecting their normalized growth scores in a two-dimensional space with the t-SNE algorithm (64). To evaluate if bacteria of the same family have a similar overall effect on cell growth relative to bacteria from other families, we applied unsupervised nearest neighbour clustering (65) and evaluated how often bacteria from the same family were nearest neighbours compared to a random sample without replacement. For this analysis, we only used bacteria from families with more than four representatives. Random groups were evaluated per family and were defined to contain the same number of strains as the family. Fisher’s exact test was used to evaluate if the nearest neighbour groups were enriched for bacteria of the same family compared to 1,000 random groups of the same size. The harmonic means of p-values from this comparison are reported.

### Association of bacterial virulence genes to growth rate changes

Bacterial genomes were screened for 35 known virulence genes that may be associated with cancer or epithelial cell changes (Supplementary Table S5). The virulence factors Map (Mitochondrial associated protein), Type-3 secretion system (T3SS) effector protein of *Citrobacter rodentium*, Shiga toxin 2A (Stx2A), Shigella enterotoxin 1 (ShEt1), and colibactin (cyclomodulin toxin on polyketide synthase (pks) island of *Enterobacteriaceae*) were identified in the *Enterobacteriaceae* family, *B. fragilis* toxin (BFT) in *B. fragilis* species, FadA in the *Fusobacteriaceae* family and *Clostridium difficile* toxin A (TcdA) in the *Clostridiales* order (Supplementary Table S5). Growth rate changes of strains containing virulence genes (if present in ≥3 strains) were compared to those of their non-virulent counterparts within the family or species (in case of *B. fragilis).* Groups were compared using Mann-Whitney U-test of aggregate data of all cell lines. P<0.05 was considered significant.

### Genes and subsystems identification

To identify genes that are associated with the inhibiting/enhancing effects of bacterial strains within families, we sorted all strains based on their effect on growth rate scores and used a normalized Kolmogorov-Smirnov statistic to quantify the association of a gene with the observed effect. Significance of the rank-based statistic was estimated by comparing its value to values obtained from 10^3^ random permutations of the same list. This provided all genes encoded by the genomes from a family with a p-value which quantified the association of the gene to the inhibiting/enhancing effect. We evaluated genes that belong to the same functional subsystems (42, 66) were more often found to be significantly associated to the inhibiting/enhancing effect than to 103 random groups of genes We defined this association by modeling the average count of genes belonging to a functional susbsystem within the random groups as Poisson distributed variables. We then tested if the count of genes with significant p-values was statistically higher than expected from the Poisson model, with a significance level of 0.05 after Benjamini/Hochberg correction for multiple tests. We used the Poisson distribution as the null distribution since (i) the labels for functional subsystem within the random groups are independent; (ii) Counts are expressed as averages from many random samples; (iii) the mean and variance are equivalent, all features consistent with a Poisson distribution(67)

## RESULTS

### Reproducibility of growth rate alterations by bacterial cells and secretomes

To identify alterations in the growth rates of CRC cell lines by bacterial cells or their secreted products, inactivated cells or secretomes were added to five CRC cell lines with different mutational landscapes and one non-cancerous kidney cell line (see Methods and Supplementary Table S2). Cell culture and bacterial culture media served as a control for cell growth for cells and secretomes respectively. A total of 7,176 experiments were conducted where we measured the bacterial effect on cell growth. Each experiment was repeated four times. We evaluated reproducibility of these replicates by performing linear regression over all pairwise combinations of replicates. As shown in Supplementary Figure S2, we found high reproducibility between replicates (regression coefficient: 0.87 ± 0.05, intercept: 0.04 ± 0.02). Based on this reproducibility, we averaged the four experimental replicates for further analysis.

### Contrasting effects between bacterial cells and secretomes

Figure 1A summarizes the effect of bacterial cells and secretomes on the growth of human cell lines. Overall, bacterial cells exhibit a stronger inhibiting effect than secretomes (p=1.02e-75, Wilcoxon signed rank test, Figure 1B). There was a low correlation between the effects of secretomes and inactivated bacteria on CRC cell growth (Supplementary Table S6), indicating that specific factors within the two compartments (cell wall attached or secreted molecules) are of importance for the observed effects.

**FIGURE 1.**
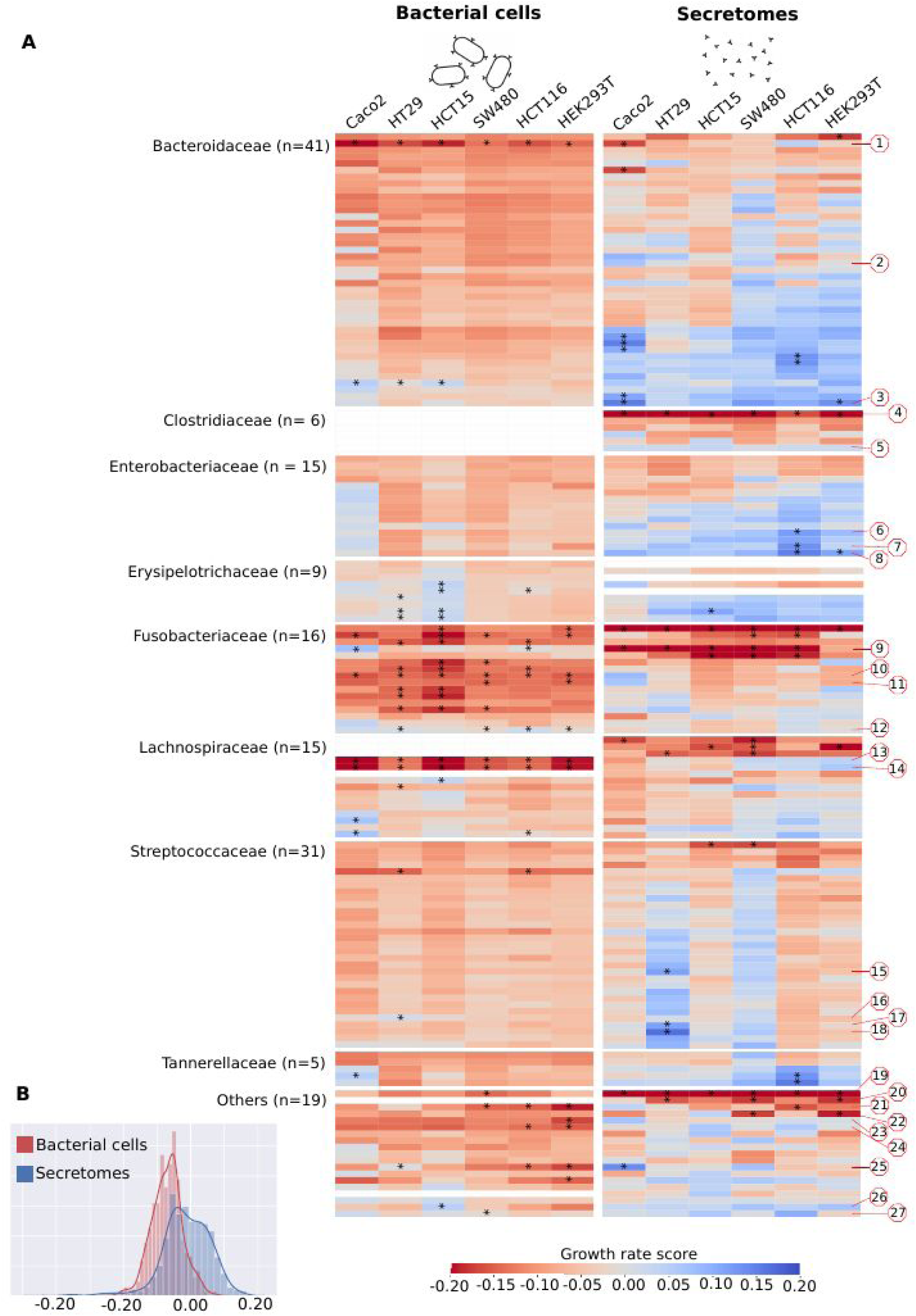
Growth rate scores of six human cell lines upon treatment with bacterial cells and secretomes. (A) Heatmap indicating low and high growth rate scores, respectively red and blue. Bacteria are sorted within bacterial families by the average growth rate score. Red numbered octagons highlight the strains discussed in the text: *Bacteroides sp.* 2_1_22 (1)*, B. fragilis* K570 clinda R (ETBF) (2), *B.* sp. 4_1_36 (3), *Clostridium septicum* (Mace 1889) Ford 1927 (4), *C.* sp. D5 (5), *Escherichia coli* D9 (6), *Klebsiella* sp. 1_1_55 (7), *E. coli* 4_1_47FAA (8), *F. nucleatum* DSM 15643 (ATCC 25586) (9), *F. nucleatum* DSM 20482 (ATCC 10953) (10), *F. necrophorum subsp. funduliforme* 1_1_36S (11), *F. nucleatum subsp. animalis* 11_3_2 (12), *Lachnospiraceae bacterium* 8_1_57FAA (13), *L. bacterium* 3_1_46FAA (14), *Streptococcus bovis* 1212 (15), *S. bovis* 1459 (16), *S. bovis* 1417 (17), *S. bovis* 207 (18), *Pediococcus acidilactici* 7_4 (19), *Pseudomonas* sp. 2_1_26 (20), *D.* sp. 6_1_46AFAA (21), *Ralstonia* sp. 5_2_56FAA (22), *Ruminococcaceae bacterium* D16 (23), *Synergistes* sp. 3_1_syn1 (24), *Desulfovibrio sp.* 3_1_syn3 (25)*, Propionibacterium sp.* 5_U_42AFAA (26), and *Eubacterium sp*. 3_1_31 (27). Highlighted with asterisks are cases that correspond to the 5% extremes of the z-score distribution. (B) Distribution of the growth rate scores for bacterial cells and secretomes.

### Difference in growth rate alterations between CRC cell lines

Growth rates were significantly enhanced or inhibited (p<0.05) in one or more cell lines in 35 out of 145 (24.1%) inactivated bacterial strains (Figure 2A, Supplementary Table S3) and 33 out of 154 (21.4%) secretomes (Figure 2B, Supplementary Table S4). Growth rates of HEK293T cells were affected by the lowest number of bacteria (n=21, n=3 enhanced and n=18 inhibited growth). On the other end of the spectrum were HCT116 cells that were affected by 28 different bacteria (n=11 enhanced and n=17 inhibited growth) and HCT15 cells that were affected by 27 different bacteria (n=8 enhanced and n=19 inhibited; Figure 2C). These two cell lines had microsatellite instability (MSI) and harbored the greatest number of pathogenic mutations in known tumor suppressor and oncogenes (n=3 and n=4 respectively, see Supplementary Table S2). In general, the number of pathogenic mutations of a cell line correlated with the number of bacteria that significantly affected its growth (Pearson r=0.931, p<0.01, see Figure 2D), suggesting that acquiring hallmark mutations might make cells more susceptible to growth rate changes by bacteria.

**FIGURE 2:**
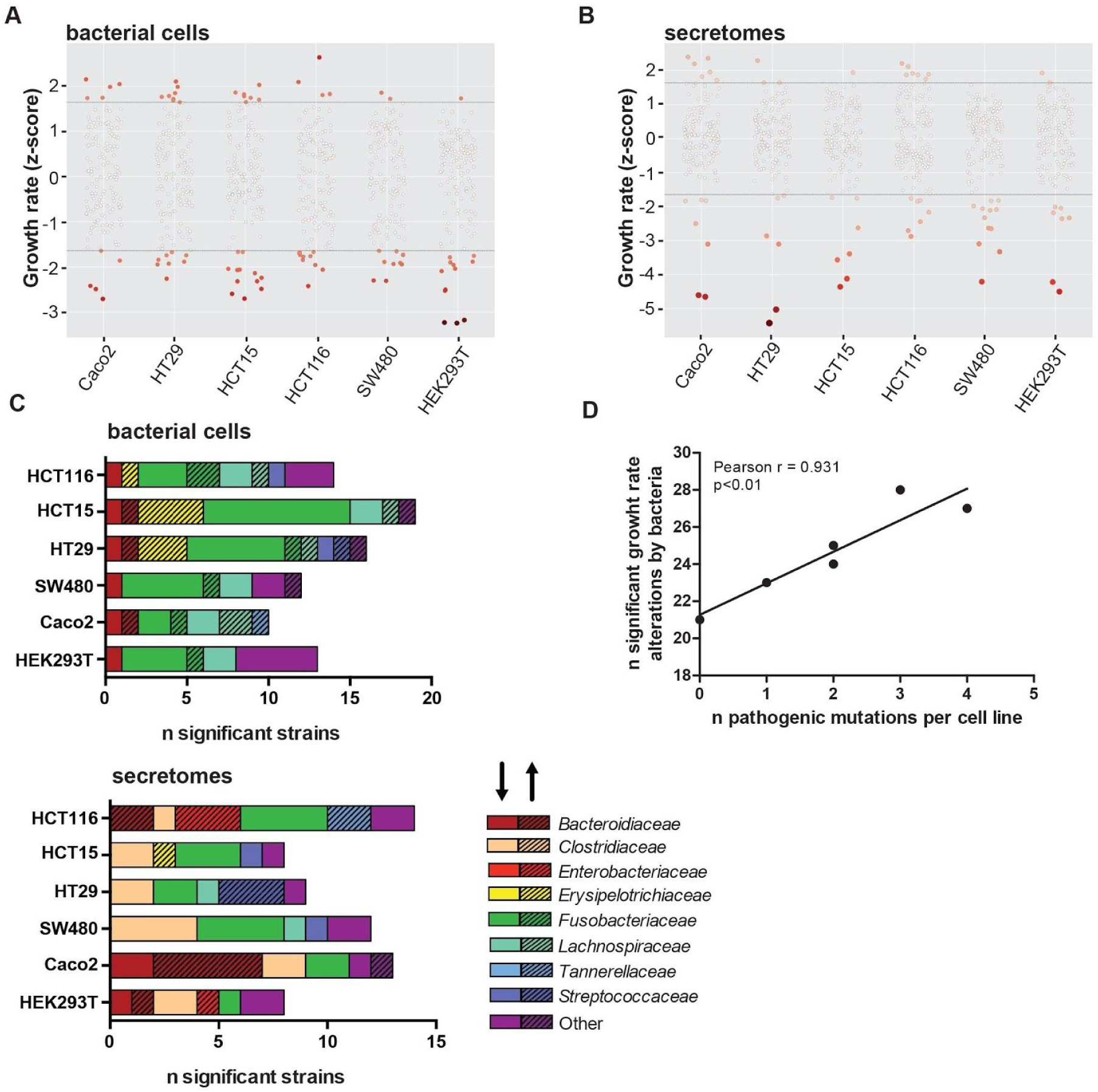
Growth rate alterations of cells lines upon incubation with bacterial cells and secretomes. Distribution plots illustrating effects of (A) bacterial cells and (B) secretomes on cell lines. Negative numbers indicate growth inhibition whereas positive numbers show growth enhancement. The color intensity of the circles represents significance of the effect on growth. (C) Overview of the number of bacterial cells and secretomes significantly enhancing or inhibiting cell growth per tested cell line, sorted by bacterial family. The shading of the bar (arrow pointing upwards) highlights enhancing and unshaded bar (arrow pointing downwards) highlights inhibiting strains. (D) Correlation of number of strains significantly altering growth rates to the number of pathogenic mutations present in cell lines. HEK293T, Caco-2, HT29, SW480, HCT116, and HCT15 were shown to possess 0, 1, 2, 2, 3, and 4 pathogenic mutations, respectively.

Some strains significantly enhanced or inhibited the growth rate of specific cell lines (Figure 1). *Streptococcaceae* cells generally had a neutral effect on cell growth, however, some *Streptococcaceae* secretomes specifically enhanced growth of HT29 and SW480 cells (Figure 1). Three *Streptococcus bovis* secretomes (strains 207, 1212 and 1417) and one inactivated bacterium (strain 1459) selectively enhanced growth in HT29 cells, while no effects were observed in other cell lines (Figure 1 and Figure 2C). The genomes of these enhancing *Streptococcus bovis* strains cluster together in a phylogenetic tree with *Streptococcus gallolyticus* subsp. *gallolyticus* (SGG) and subsp. *pasteurianus* (SGP) (Supplementary Figure 3). These effects of *Streptococcus bovis* may be cancer-specific since HEK293T was not affected. Other notable cell-line specific effects include *Erysipelotrichaceae* cells, and *Tannerellaceae* secretomes that significantly enhance growth of HCT15 and HCT116 cells, respectively.

### Bacterial family-level and strain-level effects on CRC cell growth rates

To assess the overall effect of the different bacterial secretomes and cells on our six cell lines, we projected the growth rate scores into a two-dimensional t-SNE plot. Figure 3 shows this lower dimensional space overlayed with family specific colors and a shade corresponding to the average growth effect. We observed a clear clustering of bacterial families, i.e. cells or secretomes from the same families tend to have similar effects on the different cell lines (see also Figure 1, Table 1). For bacterial cells, the most striking effect was observed for *Fusobacteriaceae* that generally inhibited growth of the cancer cells. Most *Bacteroidaceae*, *Enterobacteriaceae,* and *Erysipelotrichaceae* secretomes enhanced cell growth, although not always significantly (Figure 1). Contrastingly, *Clostridiaceae* secretomes generally inhibited growth rates. The effect of bacterial cells and secretomes was significantly clustered for five and three bacterial families, respectively (Supplementary Table S7).

**FIGURE 3.**
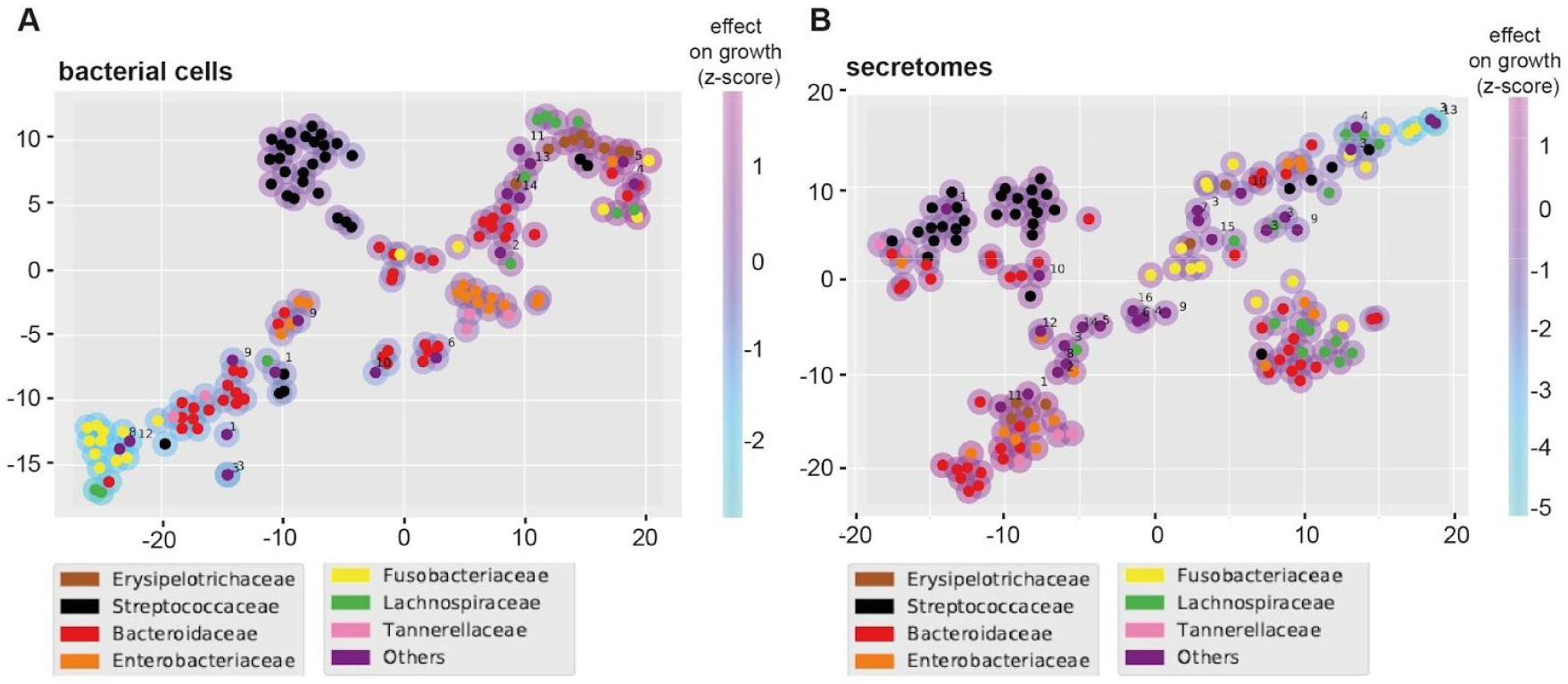
t-SNE plots summarizing the overall growth effects of bacterial cells (A) and secretomes (B) on six human cell lines. Coloured circles correspond to strains belonging to particular bacterial families. Magenta and light blue shading around the circles indicate enhancing and inhibiting effect of strains, respectively. Families labelled as “Others” contains the following strains for bacterial cells (A): *Lactobacillaceae* (1), *Peptostreptococcaceae* (2), *Desulfovibrionaceae* (3)*, Eubacteriaceae* (4), *Bifidobacteriaceae* (5)*, Enterococcaceae* (6), *Veillonellaceae* (7), *Synergistaceae* (8), *Burkholderiaceae* (9), *Pseudomonadaceae* (10), *Propionibacteriaceae* (11), *Ruminococcaceae* (12), *Akkermansiaceae* (13), *Acidaminococcaceae* (14). And the following strains for secretomes (B): *Lactobacillaceae* (1), *Eubacteriaceae* (2), *Clostridiaceae* (3), *Peptostreptococcaceae* (4), *Bifidobacteriaceae* (5)*, Enterococcaceae* (6), *Veillonellaceae* (7), *Synergistaceae* (8), *Burkholderiaceae* (9), *Desulfovibrionaceae* (10), *Propionibacteriaceae* (11), *Ruminococcaceae* (12), *Pseudomonadaceae* (13), *Akkermansiaceae* (14), *Acidaminococcaceae* (15), *Bacillaceae* (16).

**Table 1:**
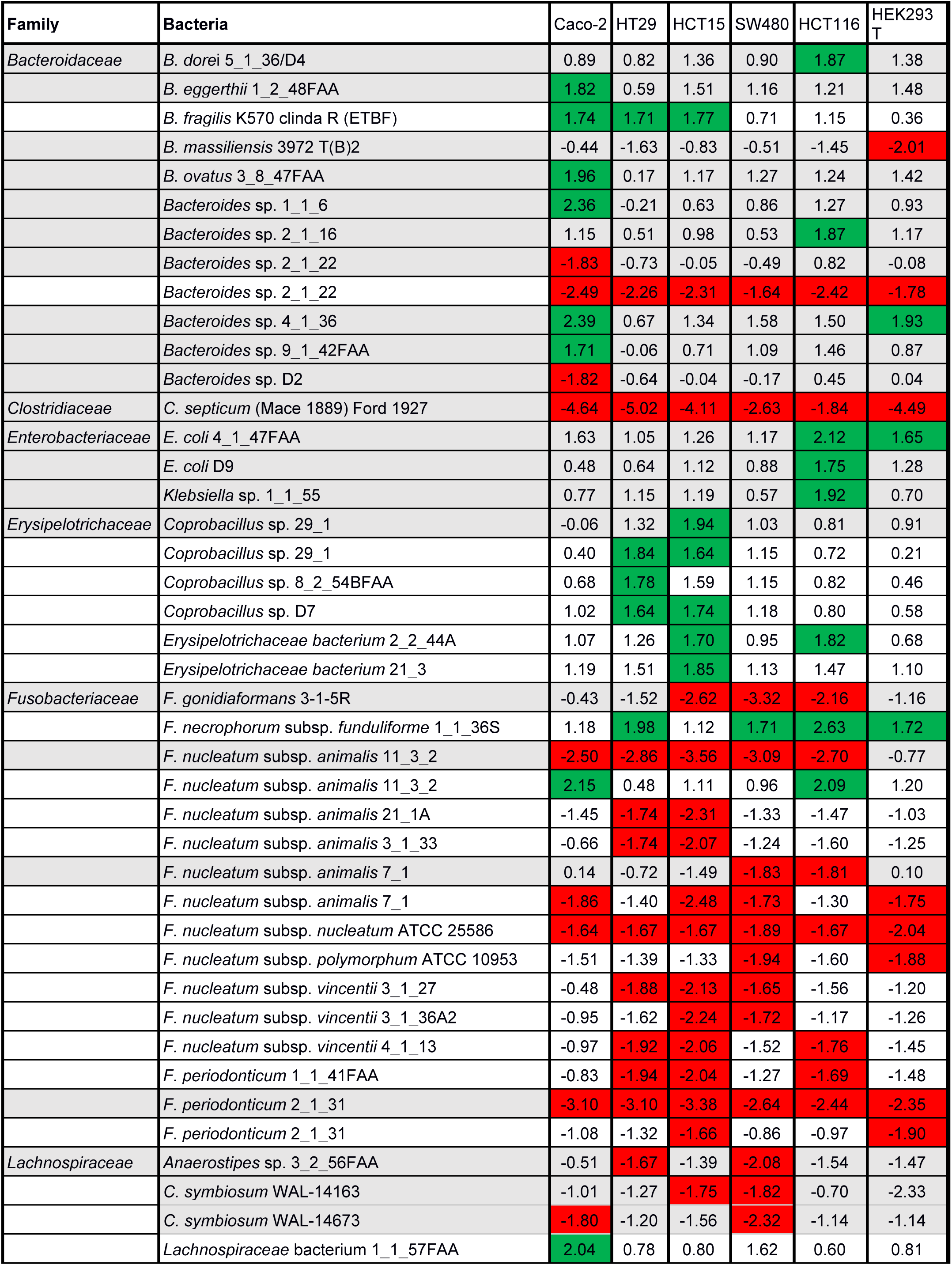
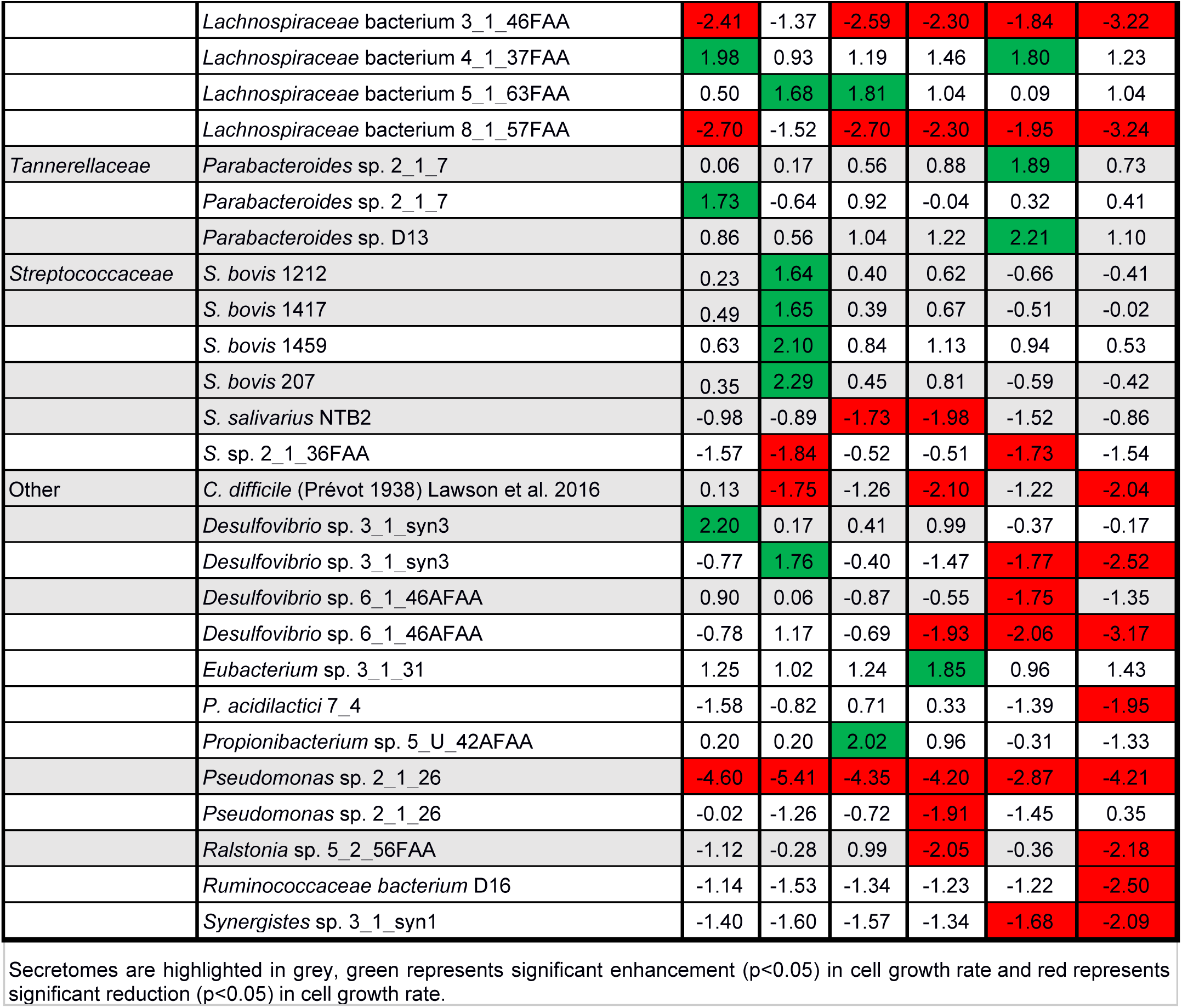
Secretomes and bacteria significantly enhancing or reducing cell growth rate (z-scores).

| Family | Bacteria | Caco-2 | HT29 | HCT15 | SW480 | HCT116 | HEK293<br>T |
| --- | --- | --- | --- | --- | --- | --- | --- |
| *Bacteroidaceae* | *B. dorei* 5_1_36/D4 | 0.89 | 0.82 | 1.36 | 0.90 | 1.87 | 1.38 |
|  | *B. eggerthii* 1_2_48FAA | 1.82 | 0.59 | 1.51 | 1.16 | 1.21 | 1.48 |
|  | *B. fragilis* K570 clinda R (ETBF) | 1.74 | 1.71 | 1.77 | 0.71 | 1.15 | 0.36 |
|  | *B. massiliensis* 3972 T(B)2 | -0.44 | -1.63 | -0.83 | -0.51 | -1.45 | -2.01 |
|  | *B. ovatus* 3_8_47FAA | 1.96 | 0.17 | 1.17 | 1.27 | 1.24 | 1.42 |
|  | *Bacteroides* sp. 1_1_6 | 2.36 | -0.21 | 0.63 | 0.86 | 1.27 | 0.93 |
|  | *Bacteroides* sp. 2_1_16 | 1.15 | 0.51 | 0.98 | 0.53 | 1.87 | 1.17 |
|  | *Bacteroides* sp. 2_1_22 | -1.83 | -0.73 | -0.05 | -0.49 | 0.82 | -0.08 |
|  | *Bacteroides* sp. 2_1_22 | -2.49 | -2.26 | -2.31 | -1.64 | -2.42 | -1.78 |
|  | *Bacteroides* sp. 4_1_36 | 2.39 | 0.67 | 1.34 | 1.58 | 1.50 | 1.93 |
|  | *Bacteroides* sp. 9_1_42FAA | 1.71 | -0.06 | 0.71 | 1.09 | 1.46 | 0.87 |
|  | *Bacteroides* sp. D2 | -1.82 | -0.64 | -0.04 | -0.17 | 0.45 | 0.04 |
| *Clostridiaceae* | *C. septicum* (Mace 1889) Ford 1927 | -4.64 | -5.02 | -4.11 | -2.63 | -1.84 | -4.49 |
| *Enterobacteriaceae* | *E. coli* 4_1_47FAA | 1.63 | 1.05 | 1.26 | 1.17 | 2.12 | 1.65 |
|  | *E. coli* D9 | 0.48 | 0.64 | 1.12 | 0.88 | 1.75 | 1.28 |
|  | *Klebsiella* sp. 1_1_55 | 0.77 | 1.15 | 1.19 | 0.57 | 1.92 | 0.70 |
| *Erysipelotrichaceae* | *Coprobacillus* sp. 29_1 | -0.06 | 1.32 | 1.94 | 1.03 | 0.81 | 0.91 |
|  | *Coprobacillus* sp. 29_1 | 0.40 | 1.84 | 1.64 | 1.15 | 0.72 | 0.21 |
|  | *Coprobacillus* sp. 8_2_54BFAA | 0.68 | 1.78 | 1.59 | 1.15 | 0.82 | 0.46 |
|  | *Coprobacillus* sp. D7 | 1.02 | 1.64 | 1.74 | 1.18 | 0.80 | 0.58 |
|  | *Erysipelotrichaceae* bacterium 2_2_44A | 1.07 | 1.26 | 1.70 | 0.95 | 1.82 | 0.68 |
|  | *Erysipelotrichaceae* bacterium 21_3 | 1.19 | 1.51 | 1.85 | 1.13 | 1.47 | 1.10 |
| *Fusobacteriaceae* | *F. gonidiaformans* 3-1-5R | -0.43 | -1.52 | -2.62 | -3.32 | -2.16 | -1.16 |
|  | *F. necrophorum* subsp. *funduliforme* 1_1_36S | 1.18 | 1.98 | 1.12 | 1.71 | 2.63 | 1.72 |
|  | *F. nucleatum* subsp. *animalis* 11_3_2 | -2.50 | -2.86 | -3.56 | -3.09 | -2.70 | -0.77 |
|  | *F. nucleatum* subsp. *animalis* 11_3_2 | 2.15 | 0.48 | 1.11 | 0.96 | 2.09 | 1.20 |
|  | *F. nucleatum* subsp. *animalis* 21_1A | -1.45 | -1.74 | -2.31 | -1.33 | -1.47 | -1.03 |
|  | *F. nucleatum* subsp. *animalis* 3_1_33 | -0.66 | -1.74 | -2.07 | -1.24 | -1.60 | -1.25 |
|  | *F. nucleatum* subsp. *animalis* 7_1 | 0.14 | -0.72 | -1.49 | -1.83 | -1.81 | 0.10 |
|  | *F. nucleatum* subsp. *animalis* 7_1 | -1.86 | -1.40 | -2.48 | -1.73 | -1.30 | -1.75 |
|  | *F. nucleatum* subsp. *nucleatum* ATCC 25586 | -1.64 | -1.67 | -1.67 | -1.89 | -1.67 | -2.04 |
|  | *F. nucleatum* subsp. *polymorphum* ATCC 10953 | -1.51 | -1.39 | -1.33 | -1.94 | -1.60 | -1.88 |
|  | *F. nucleatum* subsp. *vincentii* 3_1_27 | -0.48 | -1.88 | -2.13 | -1.65 | -1.56 | -1.20 |
|  | *F. nucleatum* subsp. *vincentii* 3_1_36A2 | -0.95 | -1.62 | -2.24 | -1.72 | -1.17 | -1.26 |
|  | *F. nucleatum* subsp. *vincentii* 4_1_13 | -0.97 | -1.92 | -2.06 | -1.52 | -1.76 | -1.45 |
|  | *F. periodonticum* 1_1_41FAA | -0.83 | -1.94 | -2.04 | -1.27 | -1.69 | -1.48 |
|  | *F. periodonticum* 2_1_31 | -3.10 | -3.10 | -3.38 | -2.64 | -2.44 | -2.35 |
|  | *F. periodonticum* 2_1_31 | -1.08 | -1.32 | -1.66 | -0.86 | -0.97 | -1.90 |
| *Lachnospiraceae* | *Anaerostipes* sp. 3_2_56FAA | -0.51 | -1.67 | -1.39 | -2.08 | -1.54 | -1.47 |
|  | *C. symbiosum* WAL-14163 | -1.01 | -1.27 | -1.75 | -1.82 | -0.70 | -2.33 |
|  | *C. symbiosum* WAL-14673 | -1.80 | -1.20 | -1.56 | -2.32 | -1.14 | -1.14 |
|  | *Lachnospiraceae* bacterium 1_1_57FAA | 2.04 | 0.78 | 0.80 | 1.62 | 0.60 | 0.81 |
|  | *Lachnospiraceae* bacterium 3_1_46FAA | -2.41 | -1.37 | -2.59 | -2.30 | -1.84 | -3.22 |
|  | *Lachnospiraceae* bacterium 4_1_37FAA | 1.98 | 0.93 | 1.19 | 1.46 | 1.80 | 1.23 |
|  | *Lachnospiraceae* bacterium 5_1_63FAA | 0.50 | 1.68 | 1.81 | 1.04 | 0.09 | 1.04 |
|  | *Lachnospiraceae* bacterium 8_1_57FAA | -2.70 | -1.52 | -2.70 | -2.30 | -1.95 | -3.24 |
| *Tannerellaceae* | *Parabacteroides* sp. 2_1_7 | 0.06 | 0.17 | 0.56 | 0.88 | 1.89 | 0.73 |
|  | *Parabacteroides* sp. 2_1_7 | 1.73 | -0.64 | 0.92 | -0.04 | 0.32 | 0.41 |
|  | *Parabacteroides* sp. D13 | 0.86 | 0.56 | 1.04 | 1.22 | 2.21 | 1.10 |
| *Streptococcaceae* | *S. bovis* 1212 | 0.23 | 1.64 | 0.40 | 0.62 | -0.66 | -0.41 |
|  | *S. bovis* 1417 | 0.49 | 1.65 | 0.39 | 0.67 | -0.51 | -0.02 |
|  | *S. bovis* 1459 | 0.63 | 2.10 | 0.84 | 1.13 | 0.94 | 0.53 |
|  | *S. bovis* 207 | 0.35 | 2.29 | 0.45 | 0.81 | -0.59 | -0.42 |
|  | *S. salivarius* NTB2 | -0.98 | -0.89 | -1.73 | -1.98 | -1.52 | -0.86 |
|  | *S. sp.* 2_1_36FAA | -1.57 | -1.84 | -0.52 | -0.51 | -1.73 | -1.54 |
| Other | *C. difficile* (Prévot 1938) Lawson et al. 2016 | 0.13 | -1.75 | -1.26 | -2.10 | -1.22 | -2.04 |
|  | *Desulfovibrio* sp. 3_1_syn3 | 2.20 | 0.17 | 0.41 | 0.99 | -0.37 | -0.17 |
|  | *Desulfovibrio* sp. 3_1_syn3 | -0.77 | 1.76 | -0.40 | -1.47 | -1.77 | -2.52 |
|  | *Desulfovibrio* sp. 6_1_46AFAA | 0.90 | 0.06 | -0.87 | -0.55 | -1.75 | -1.35 |
|  | *Desulfovibrio* sp. 6_1_46AFAA | -0.78 | 1.17 | -0.69 | -1.93 | -2.06 | -3.17 |
|  | *Eubacterium* sp. 3_1_31 | 1.25 | 1.02 | 1.24 | 1.85 | 0.96 | 1.43 |
|  | *P. acidilactici* 7_4 | -1.58 | -0.82 | 0.71 | 0.33 | -1.39 | -1.95 |
|  | *Propionibacterium* sp. 5_U_42AFAA | 0.20 | 0.20 | 2.02 | 0.96 | -0.31 | -1.33 |
|  | *Pseudomonas* sp. 2_1_26 | -4.60 | -5.41 | -4.35 | -4.20 | -2.87 | -4.21 |
|  | *Pseudomonas* sp. 2_1_26 | -0.02 | -1.26 | -0.72 | -1.91 | -1.45 | 0.35 |
|  | *Ralstonia* sp. 5_2_56FAA | -1.12 | -0.28 | 0.99 | -2.05 | -0.36 | -2.18 |
|  | *Ruminococcaceae* bacterium D16 | -1.14 | -1.53 | -1.34 | -1.23 | -1.22 | -2.50 |
|  | *Synergistes* sp. 3_1_syn1 | -1.40 | -1.60 | -1.57 | -1.34 | -1.68 | -2.09 |

Secretomes are highlighted in grey, green represents significant enhancement ( $p < 0.05$ ) in cell growth rate and red represents significant reduction ( $p < 0.05$ ) in cell growth rate.

In the *Bacteroidaceae* family (n=41 strains tested), secretomes and bacterial cells of seven and one strains significantly enhanced cell growth, respectively. The effects were most prominently observed in Caco-2 cells, while the growth of non-cancerous cell line HEK293T was only significantly enhanced by the secretome of *Bacteroides* sp. 4_1_36 (Figure 2C & Table 1). Only three *Enterobacteriaceae* secretomes (n=15) significantly enhanced growth of HEK293T cells (*Escherichia coli* 4_1_47FAA) and HCT116 (*Escherichia coli* 4_1_47FAA and D9, and *Klebsiella* sp. 1_1_55) while no effects were observed for bacterial cells.

Growth inhibiting effects were mainly observed within the family *Fusobacteriaceae* (Figure 1, Table 1). From the 12 out of 16 significant *Fusobacteriaceae* (81.3%) cells, two enhanced and ten inhibited cellular growth. *Fusobacterium necrophorum* subsp. *funduliforme* 1_1_36S cells enhanced growth in HCT116, SW480, and HEK293T cells. Strikingly, while *Fusobacterium nucleatum* subsp. *animalis* 11_3_2 cells enhanced growth of Caco-2 and HCT116, its secretome inhibited growth in those same cell lines. Especially the cell line HCT15 was sensitive for *Fusobacteriaceae* growth inhibition where 11 out of 16 strains significantly reduced growth rates.

Alternatively, HEK293T and Caco-2 were less sensitive to *Fusobacteriaceae* with only four out of 16 strains inhibiting cell growth. Notably, the effect of *Fusobacteriaceae* secretomes on cell growth may be cancer specific, since most secretomes that inhibited cancer cell growth did not inhibit growth of HEK293T cells.

### Families with fewer than 5 strains tested

Among strains from families with less than five tested strains (n=18), *Eubacterium sp.* 3_1_31 and *Propionibacterium sp.* 5_U_42AFAA cells increased cell growth rates in SW480 and HCT15 cells, respectively (Table 1). Interestingly, *Pseudomonas* sp. 2_1_26 secretomes significantly inhibited the growth rates of all cell lines while the cells specifically inhibited SW480. *Pediococcus acidilactici* 7_4 and *Ruminococcaceae bacterium* D16 cells selectively inhibited the non-cancerous cell line, HEK293T, while *Synergistes sp.* 3_1_syn1 also inhibited HCT116. Aerobically cultured *Ralstonia* sp. 5_2_56FAA secretomes inhibited 2 cell lines, SW480 and HEK293T. For *Desulfovibrio* sp. 3_1_syn3, the secretome increased Caco-2 growth while the bacterial cells increased HT29 growth. Contrastingly, these same bacterial cells decreased growth of HCT116 and HEK293T. Similarly, *Desulfovibrio* sp. 6_1_46AFAA cells decreased the growth rates of HCT116, SW480 and HEK293T cells.

### Strain-specific effects that contrast their family

While strains from the same bacterial families tend to have similar effects on the growth rates of cell lines (Supplementary Table S7), large variations were observed within certain families. For instance, within the *Clostridiaceae* family, the secretome of *Clostridium septicum* (Mace 1889) Ford 1927 strongly inhibited growth of all tested cell lines, while the *Clostridium* sp. D5 secretome enhanced growth, although the significance threshold was not reached (Figure 1 and Supplementary Table S4). Similarly, *Bacteroides* sp. 2_1_22 cells strongly inhibited the growth rates of all cell lines while *B. fragilis* K570 cells enhanced growth in most cell lines (Figure 1 and Supplementary Table S3). A large growth rate variation was also observed in cell lines exposed to *Lachnospiraceae* cells (n=11). For this family, the cells of *L. bacterium* 3_1_46FAA and *L. bacterium* 8_1_57FAA significantly inhibited the growth rates of all cell lines, while most of the other strains showed weak or low inhibition.

The observed variations in growth effect within bacterial families might either be explained by the phylogenetic distances between the strains or by characteristics with a non-phylogenetic distribution, such as accessory functions. To evaluate this, we measured the distances on a phylogenomic tree and compared them with the differences in growth rate effects over the cell lines. For most bacterial families, there is no association between the growth rate effects and phylogenetic distances (Supplementary Table S8, Supplementary Figure S4), suggesting that these effects may be associated with accessory functions that are independently gained or lost in different lineages. For *Erysipelotrichaceae* secretomes and cells, *Fusobacteriaceae* cells, and *Lachnospiraceae* secretomes, we found that the phylogenetic distance was significantly correlated with the growth rate effects, suggesting that these effects may be associated to variations in universally shared genes or to accessory functions with phylogenetically consistent dynamics.

### The presence of virulence genes is associated with growth rate inhibition

To explain the complex, sometimes strain-dependent patterns of growth enhancement or inhibition, we analyzed the genome sequences of 144/157 tested gut bacteria. First, we checked whether 35 known bacterial family-specific virulence genes were present within the genomes of the tested strains (Supplementary Table S9), because some of the encoded virulence factors have been linked to cell proliferation changes. Seven toxin genes were found, including TcdA (68); FadA (31); colibactin present on the pks-island (69, 70); Shiga toxin ShEt1; *Shigella* enterotoxin Stx2A; effector protein Map secreted by T3SS of *Citrobacter rodentium* and *Escherichia coli* (71, 72); and BFT (38).

As shown in Figure 4, strains containing virulence genes tend to change growth rates, relative to related strains without those virulence genes. For example, the secretomes of *Clostridiales* strains significantly inhibited growth rate if their genomes contained *TcdA* (Figure 4A, p<0.0001). Similarly, *Fusobacteria* strains encoding the membrane protein *fadA*, that might be responsible for growth rate changes of epithelial cells (31), inhibited growth rates significantly (Figure 4B, p<0.0001 and p<0.05 for bacterial cells and secretomes respectively). Within the *Enterobacteriaceae* family, multiple secreted toxins were identified in the genomes, including colibactin, *ShEt1*, *Stx2A*, and *Map*. Secretomes of *Enterobacteriaceae* possessing any of these toxins inhibited cell growth relative to strains without these toxins (Figure 4C, p<0.0001). No differences were observed between bacterial cells of *Enterobacteriaceae* with or without the toxin genes. The secretomes of three *B. fragilis* strains encoding the secreted enterotoxin *bft* enhanced the growth rates of CRC cells, while the five *B. fragilis* strains without the toxin did not, although this trend was not significant (Figure 4A, p=0.075). The trend was not observed for bacterial cells, where only *B. fragilis* K570 significantly enhanced growth rates (Table 1).

**FIGURE 4.**
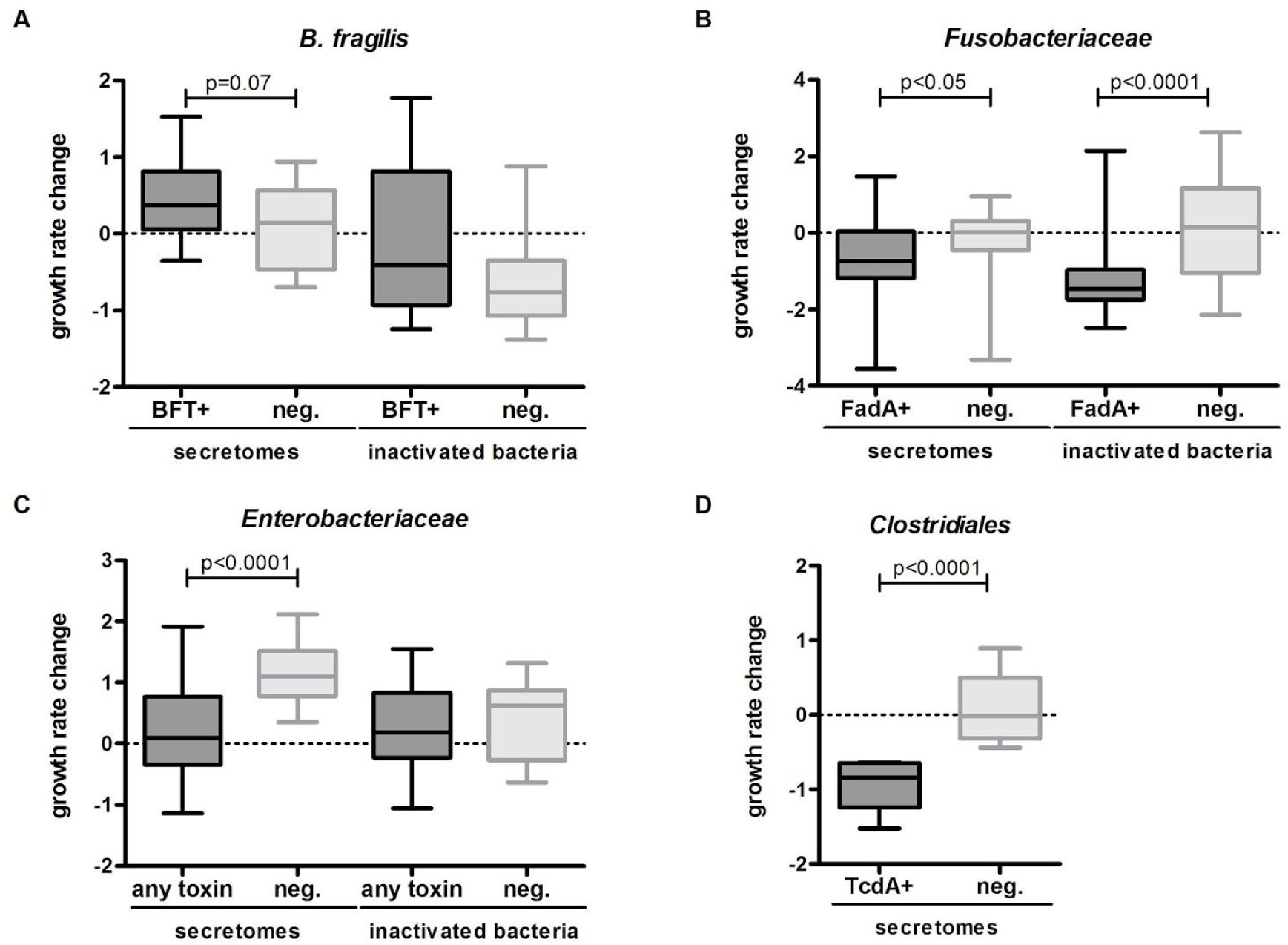
Influence of strains with and without encoded virulence factors on cellular growth rates. *B. fragilis* (A), *Fusobacteriaceae* (B), *Enterobacteriaceae* (C) and *Clostridiaceae* (D) encoding the virulence factors *bft*, *fadA*, any toxin (*Map*, *pks*, *ShEt1*, *Stx2A*), and *TcdA*, respectively, were compared to strains without these encoded virulence factors (Mann-Whitney U test). A trend towards cell growth enhancement was observed for *B. fragilis bft*+ secretomes (p=0.07) (A), while significant inhibition of cell growth was observed for *fadA*+ *Fusobacteriaceae* cells (p<0.0001), and secretomes (p<0.05) (B), for *Enterobacteriaceae* secretomes encoding toxins (p<0.0001) (C), and *TcdA*+ *Clostridiales* secretomes (p<0.0001) (D).

### Functional categories associated with differential effects within families

Beside the toxin genes described above, other genes might also play a role in the observed effects. Thus, we next performed an unbiased comparative genome analysis where we associated all annotated bacterial genes to the effect of the strain on cell growth. We next grouped the genes by functional category and assessed whether genes within a category were significantly associated with cell growth alterations. Based on the PATRIC (42) annotations we evaluated genes in two hierarchical levels of functions, namely ‘Subsystems’ and ‘Superclass’. All functional categories found to be significantly associated with cell growth alterations are listed in Supplementary Table S10.

The “Membrane transport” superclass was significantly associated with inhibiting *Enterobacteriaceae* cells and secretomes, and with enhancing *Fusobacteriaceae* secretomes. Other enriched superclasses included: “Cell envelope”, “Metabolism”, “Cellular processes”, “Protein processing”, “Energy”, and “Stress response, defense and virulence” (Supplementary Table S10). Subsystems including “Cobalamin synthesis”, “Ethanolamine utilization”, and the “Dpp dipeptide ABC transport system” were consistently found to be associated with growth enhancement or inhibition in multiple bacterial families and in multiple cell lines (Supplementary Table S10). The significant effects of other subsystems including “Vir-like type 4 secretion system”, “Glutamate fermentation”, “Flagellum”, “Branched-chain amino acid biosynthesis”, and others were associated with specific bacterial families. We expect that these statistical associations will provide valuable leads for future experiments.

## DISCUSSION

Over 150 human gut bacteria belonging to taxonomic groups that have previously been associated with CRC were tested for their effects on cell growth of five CRC cell lines (HT29, Caco-2, HCT116, SW480 and HCT15) with different mutation landscapes representing commonly mutated genes in CRC, and one non-cancerous kidney cell line (HEK293T). Our results show that cell growth enhancing and inhibiting effects of gut bacteria depend both on the mutational landscape of the cancer cells and on the identity and functional content of the bacteria, including secreted or cell wall attached factors. CRC is a heterogeneous disease, as is reflected in the five different CRC cell lines with mutations in key CRC related genes (*KRAS*, *APC, BRAF*, *TP53*, *CTNNB1* and *PIK3CA*). As expected, the growth rate of non-cancerous HEK293T kidney cells was affected by the lowest number of gut bacterial strains, with only small differences.

### Closely related bacteria may have distinct effects on cell growth

A striking observation was that the effects of bacterial cells and secretomes were strongly associated to bacterial families, although exceptions to this general pattern were observed. Previous studies have found that *Fusobacteria*, *Streptococcus, Peptostreptococcus, Ruminococcus* and *Escherichia/Shigella* tend to be enriched in tumors, while *Bacteroides, Pseudomonas, Lachnospira* and *Anaerostipes* are depleted (5, 7, 9, 22–24, 28, 31, 41, 73–76). Regardless of the enrichment or depletion on CRC tumors *in vivo,* our *in vitro* results indicate that closely related bacteria may have distinct effects on cancer cell growth. For example, we observed that different *Fusobacteria* strains, which were all cultured in the same media under the same conditions, may have opposing effects on cancer cell growth, while these bacteria are consistently enriched in patients (24, 28, 77). This suggests that specific bacterial factors play a role in CRC.

### *Fusobacteriaceae*: enhancing or inhibiting growth?

Previous studies indicated that *Fusobacteriaceae* enhanced growth of HCT116, SW480, and HT29 cells (31, 76). *F. nucleatum* ATCC25586was shown to enhance CRC cell growth via interaction with Toll like receptors TLR4, TLR2, and myeloid differentiation primary response protein 88 (MYD88) (76), and the *F. nucleatum* strain Fn12230 was previously shown to enhance cell growth through the cell-wall attached adhesin FadA (31). Our results showed that most *Fusobacteriaceae* significantly inhibited cellular growth, including *Fusobacterium nucleatum* ATCC25586, contrasting previous observations (76, 78), and that the presence of the *FadA* gene was significantly linked to growth inhibition by cells and secretomes (Table 1). Notably, the secretomes showed less pronounced inhibition than bacterial cells. While FadA is membrane-bound, secretomes likely include outer membrane vesicles (OMVs) that are 40-110 nm in size and express FadA (32, 79). To rule out the possibility that these contrasting results were due to differences in methods, we compared our high-throughput MTT-based assay, used in this study, with the cell counting assay, that was employed in the referred studies, on 6 cell lines, using 6 different secretomes, and found a high correlation (Pearson’s r=0.813, Supplementary Figure S5). These results confirm that the observed inhibitory effects by *Fusobacteriaceae* strains do not depend on the applied experimental techniques, but may partly be due to use of different *Fusobacteriaceae* strains(Fn12230, ATCC25586, ATCC23726) isolated from extra-intestinal sites(80). Specifically HCT15 cell growth was significantly inhibited by 11/16 *Fusobacteriaceae* strains (nine cells and three secretomes). HCT15 cells contain a missense mutation in MYD88 (p.Arg264Ter) resulting in a MYD88 deficiency, and may thus explain the insensitivity of this cell line to *F. nucleatum* subsp. *animalis* 11_3_2 and *F. necrophorum* subsp. *funduliforme* 1_1_36S, the two *Fusobacteriaceae* strains that promoted growth of the other CRC cell lines, but not of HCT15.

### Virulence factors are associated to cell growth inhibition

The growth-inhibiting properties of *Clostridiaceae* secretomes may result from toxins encoded by this family. We observed that the presence of the *TcdA* gene in *Clostridiaceae* was significantly associated with cell growth inhibition. *Clostridium septicum* (Mace 1889) Ford 1927 secretomes inhibited cell growth in all of our cell lines. This strain produces multiple exotoxins (alpha toxin, lethal toxin and hemolytic toxin) and has been associated to CRC (14, 15).

*Enterobacteriaceae* strains encoding the toxins colibactin, *Stx2A*, *ShEt1*, and *Map* inhibited cell growth relative to non-toxigenic *Enterobacteriaceae* strains. Colibactin (present in four of our *Escherichia* strains) is a genotoxic compound that alkylates DNA and generates DNA double strand breaks (41, 74). The DNA damaging effect of colibactin may not directly affect cell growth in our experimental setup, although colibactin may induce cell cycle arrest via SUMOylation of TP53 causing inactivation (69, 70). Cell cycle arrest would stall cell growth, which can only happen in *TP53* wildtype cell lines with active TP53 protein. Indeed, the only cell lines that were relatively inhibited by strains encoding colibactin were the wildtype *TP53* cell lines HEK293T and HCT116.

An important virulence factor of *B. fragilis* is BFT (81). While in previous research BFT was shown to enhance cell growth of HT29/c1 cells via β-catenin nuclear signalling (38), enterotoxigenic *B. fragilis* secretomes (VPI13784 (*bft1*), 86-5443-2-2 (*bft2),* and K570 (*bft3*)) only marginally enhanced CRC cell growth in our assay compared to the other *B. fragilis* secretomes in our screen. The mild effects as observed may depend on the level of toxin production and secretion under our experimental conditions. Interestingly, enterotoxigenic *B. fragilis* str. Korea 570 cells significantly enhanced growth, which might be independent of BFT and related to other cell-wall attached factors.

### *Streptococcus bovis* strains specifically enhance growth of HT29 cells

*Streptococcus bovis* infections have been linked to colorectal neoplasia (43, 82, 83). Our results demonstrate that secretomes of *Streptococcus bovis* strains that are closely related to known *S. gallolyticus* and *S. pasteurianus* strains (Supplementary Figure S3), selectively enhance growth of HT29 cells. This is in line with a previous study showing cell growth enhancing effects in HT29 and HCT116 cells that was mediated through *β-*catenin, while no effect was observed in SW480 and HEK293T cells (84). An intact *β-*catenin pathway may therefore be required to observe the specific effect of *S. gallolyticus* on cell growth as is the case in HT29 cells (85), which may thus be sensitive to the effects of *S. gallolyticus* in our assay. HCT116 cells have a mutation in *CTNNB1 (p.Ser45del)* that interferes with *β-*catenin degradation, which may explain its insensitivity to *S. gallolyticus*. Alternatively, mutations in the *APC*-gene in the other CRC-cells (86) may interfere with *β-*catenin signalling depending on the specific mutation.

### Bacterial functions associated with cell growth alterations

Several functional categories were significantly associated with CRC cell growth. Most of these functions were identified in specific bacterial families and consistently inhibited or enhanced cell growth in different cell lines. These functions may reflect novel pathways of bacterial interference with CRC cell growth that, to our knowledge, have not been previously identified. Most functions are related to cell metabolism, secretion, and transport systems. For example, secretomes of *Enterobacteriaceae* that encode the “Vir-like type 4 secretion system” inhibited growth of all cell lines. Similarly, the gene superclass “Membrane transport” was mostly associated to secretomes that inhibited cell growth (Supplementary Table S10). Molecules that are secreted by these bacterial transport systems may be responsible for the inhibiting effect. For example, the Vir-like type 4 secretion system allows bacteria to secrete proteinaceous effectors that kill competitors (87). Here, we report an important first step in understanding the cell growth enhancing or inhibiting effects of human gut bacteria by identifying putative transport systems for effector molecules.

While the growth effects of secretomes were mostly associated with membrane transport functions, effects on CRC cell growth by bacterial cells were associated with metabolic pathways (Supplementary Table S10). It is possible that these associations may be attributed to metabolic enzymes that may still be, although the bacterial cells were inactivated. In other ecosystems, metabolic enzymes of dead bacteria have been shown to remain active for up to 96 hours (88). Our experiment was performed within 72 hours. The synthesis of cobalamin (vitamin B12), which is important for human cell proliferation *in vitro* (89) and a common addition to cell culturing media, was significantly associated with the cell growth enhancing effect of *Bacteroidaceae* and *Enterobacteriaceae* cells. Several possible scenarios can be envisioned to explain this observation. First, it remains to be tested whether vitamin B12 was retained in the bacterial fraction after washing, stimulating human cell growth on those cells. Second, different human gut-associated *Bacteroides* strains contain surface-exposed lipoproteins with high affinity for cobalamin (90). Such molecules may remove cobalamin directly from the media, making it unavailable to the human cells and impeding their growth.

The “Ethanolamine utilization” pathway was another metabolic process that we found to be associated to the inhibiting effect of *Fusobacteriaceae* cells on the growth rate of CRC cell lines. Ethanolamine (EA) is the basal component of phosphatidylethanolamine, a major phospholipid in animal cell membranes and is utilized by bacteria including *Fusobacteria* as a carbon source in the gut (91–93). EA sensing regulators have been proposed to regulate virulence in *Enterobacteriaceae* (93, 94). It remains to be tested if the EA in human cell membranes triggers virulence factors in *Fusobacteria* that encode EA utilization genes, or perhaps if inactivated *Fusobacteria* cells retain the capacity to digest and kill human cells by the activity of EA utilization enzymes.

Overall, the statistical associations found in this exploratory analysis provide valuable clues about the possible functions and mechanisms that drive the interactions of bacteria with human cells in the gut. Future experiments are needed to confirm whether these factors are causal and to further clarify the molecular mechanisms involved.

### Outlook

Our broad screen revealed many significant associations between gut bacteria, their encoded functions, and growth enhancement or inhibition of human CRC cell lines. Many growth rate effects remain elusive and need to be further examined. Moreover, it will be important to investigate the growth rate effects of living bacterial communities or mock communities, which are thought to synergize in affecting tumorigenesis (53), but fall outside the scope of our present study. Bacterial effects should be examined in more natural environments that mimic the conditions in the human gut, *e.g.* in co-culture with mucus, immune cells, and/or within organoid cultures (95). Our study forms an important baseline for such further research by identifying important traits in individual bacterial strains that are associated to enhancing or inhibiting cancer cell growth, as well as important host factors that determine sensitivity to those bacterial traits.

## CONCLUSIONS

Above, we presented the results of the first large-scale analysis of the effects of gut bacteria on CRC cell growth. Our results with bacterial cells and secretomes reflect the complexity of the microbe-host interactions in the human gut. First, bacterial families tend to have consistent effects on cell growth, although these effects are not universal. This may partially be explained by the presence of virulence genes in some strains. Second, there are cell line dependent effects in CRC cells due to their mutational landscape heterogeneity. Notably, these cell lines have acquired several different cancer hallmarks, yet their growth can still be influenced by specific bacteria. Our results suggest that the response of CRC cells to gut bacteria may differ between patients due to differences in their gut bacteria, although the relevance of our *in vitro* results for patients remains to be studied. Cell growth enhancing and inhibiting bacterial traits could be important for determining risk for cancer or its progression, and these consequences could potentially be counteracted by microbiome modulation. Currently, microbiome medicine applications have already been adapted to some other diseases. For instance, *Akkermansia muciniphila* was demonstrated to improve diabetes type 2 and obesity in mice (17, 18). Resistant life-threatening *Clostridium difficile* infections and ulcerative colitis were successfully treated with faecal microbiota transplants (FMT) (19, 20). Our study highlights the complex processes at play at the gut-microbiome interface and contributes to a better understanding of the role of different bacterial strains in altering colonic epithelial cell proliferation.

## Acknowledgements

We thank Shaoguang Wu and Cynthia Sears from Johns Hopkins Medical Institutions for providing different gut bacterial strains (see Methods).

## Supplementary Figure captions

**S1.**
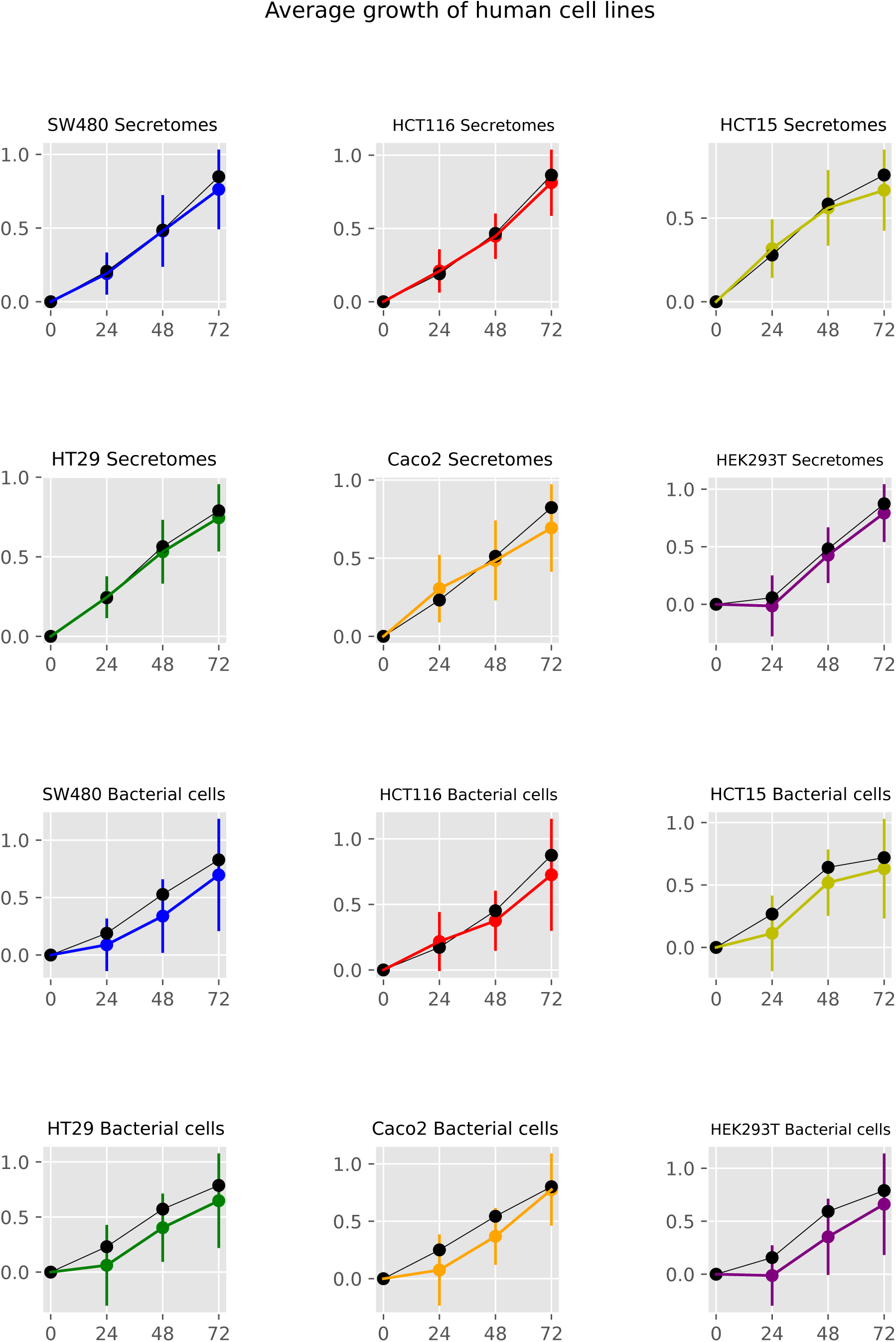
Average growth of human cell lines exposed to bacterial cells and secretomes. Colored lines and error bars show the average and standard deviation of MTT measurements for 6 human cell lines exposed to cells and secretomes of 145 and 154 bacteria, respectively. Measurements were recorded at the incubation times of 0, 24, 48, and 72h. The dark lines show the average of the cell line without bacterial products (negative controls).

**S2.**
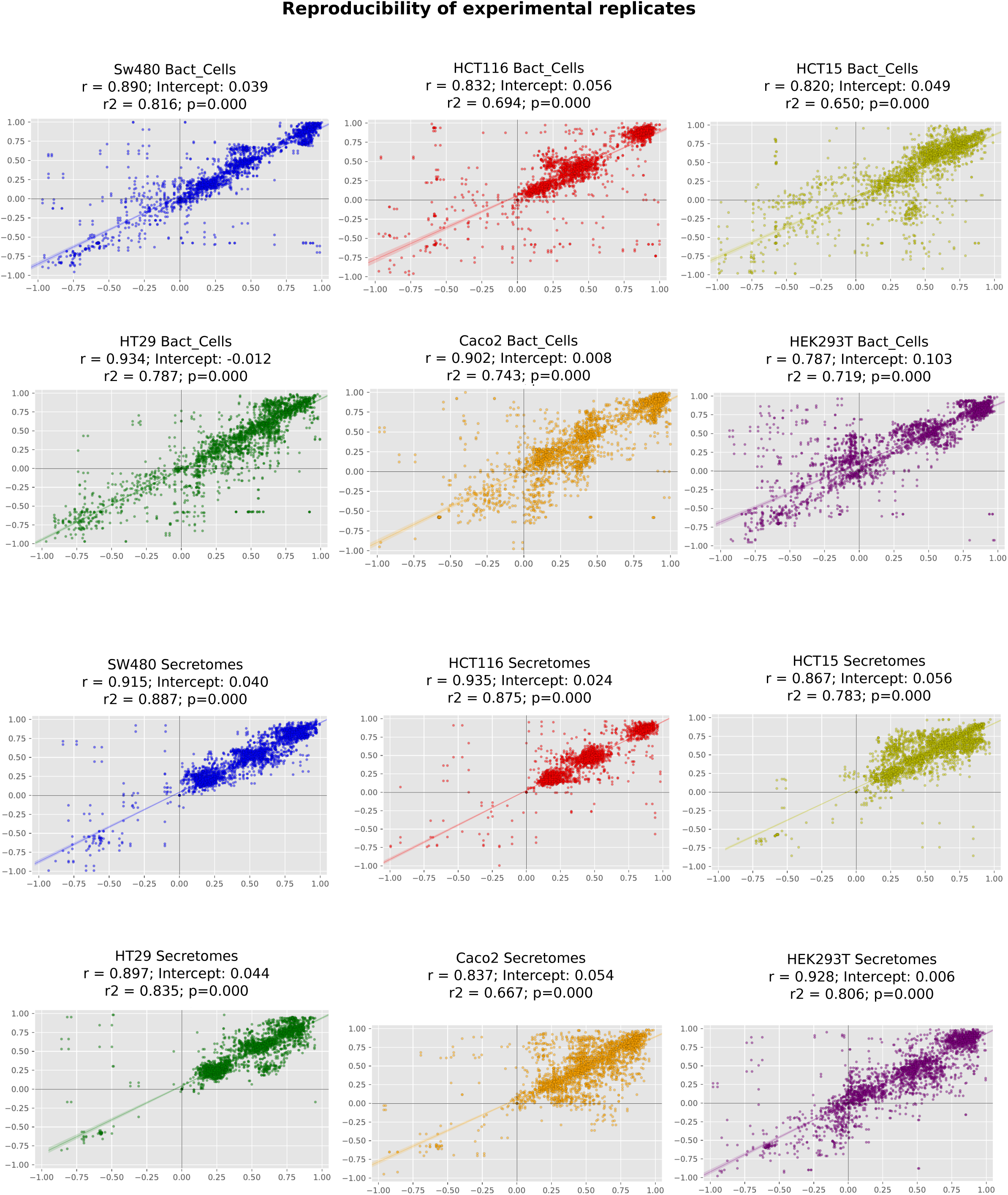
Reproducibility of four experimental replicates. Scatter plots of the pairwise combinations of four experimental replicates of MTT measurements at incubation times of 0, 24, 48, and 72h. Individual plots exhibit replicates of one cell line exposed to the cells or secretomes of, respectively 145 and 154 bacterial strains. Lines show the best fit to a linear model. The specific linear regression coefficients, intercepts, R^2^, and p-values are shown on the top of each plot.

**S3.**
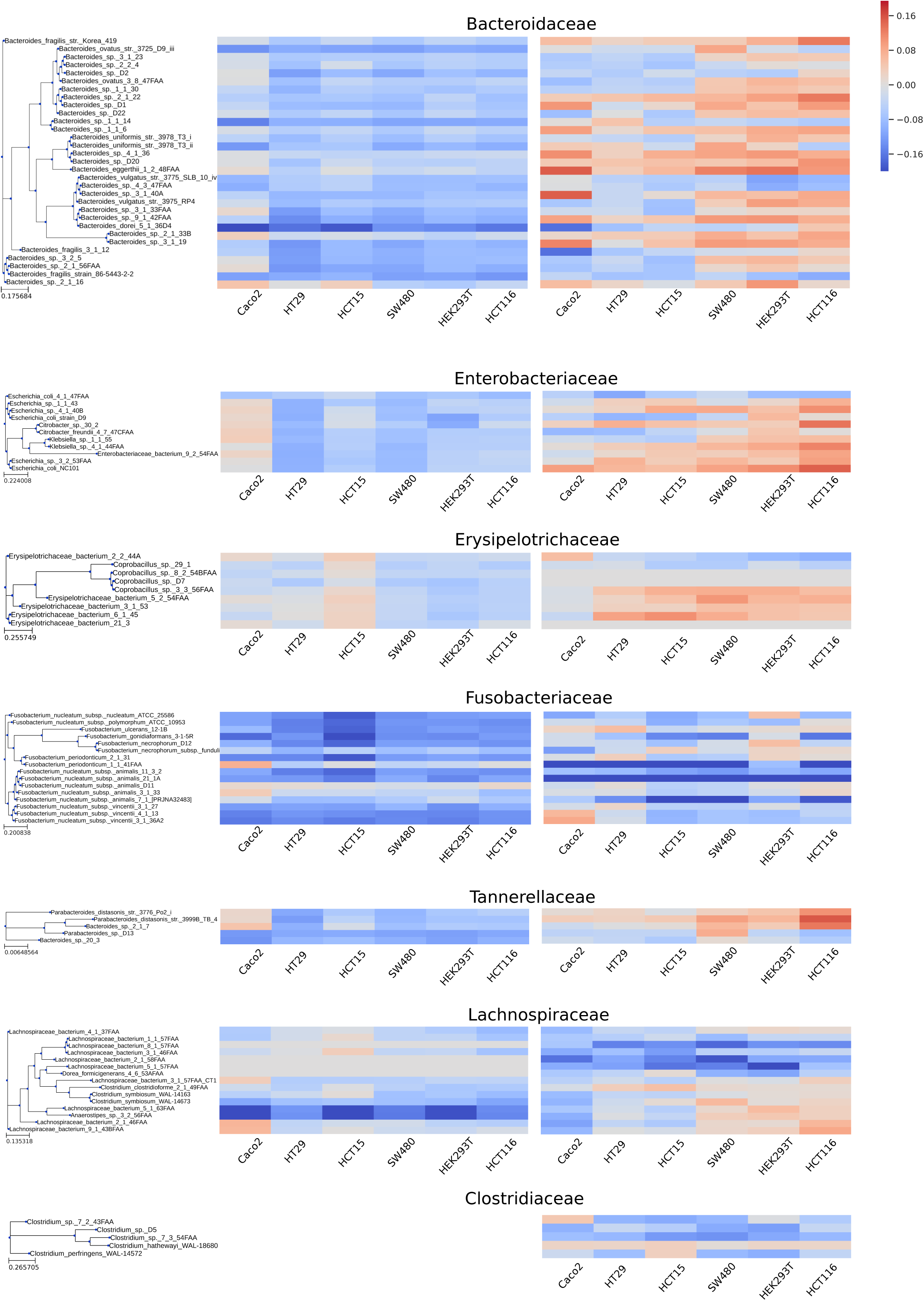
Cell growth rate alterations ordered by phylogenomic distances. Heatmaps show the growth rate alterations of six cell lines incubated with the cells and secretomes of strains from seven bacterial families. Rows of the heatmap are ranked according to the tips of the phylogenomic trees shown on the left.

**S4.**
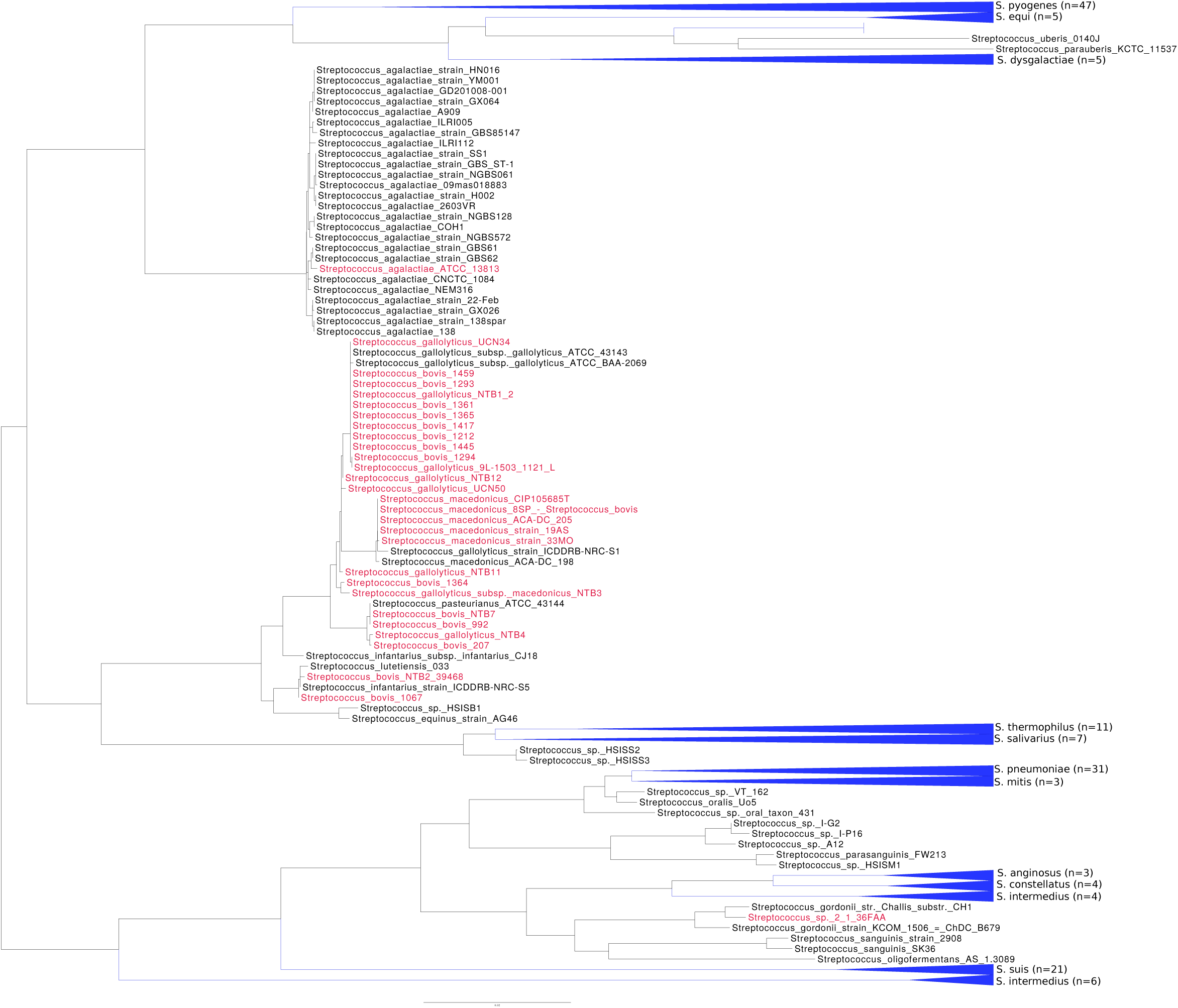
*Streptococcus* phylogenomic tree. Genome-based phylogenetic reconstruction of 238 Streptococcus genomes confirms that the genome sequences of the *S. bovis* strains reported in this study are closely related to previously sequenced genomes of *S. gallolyticus* and *S. pasteurianus*. The taxa names of strains used in our study are highlighted red. Monophyletic branches containing strains from a single species are collapsed and highlighted with blue triangles.

**S5.**
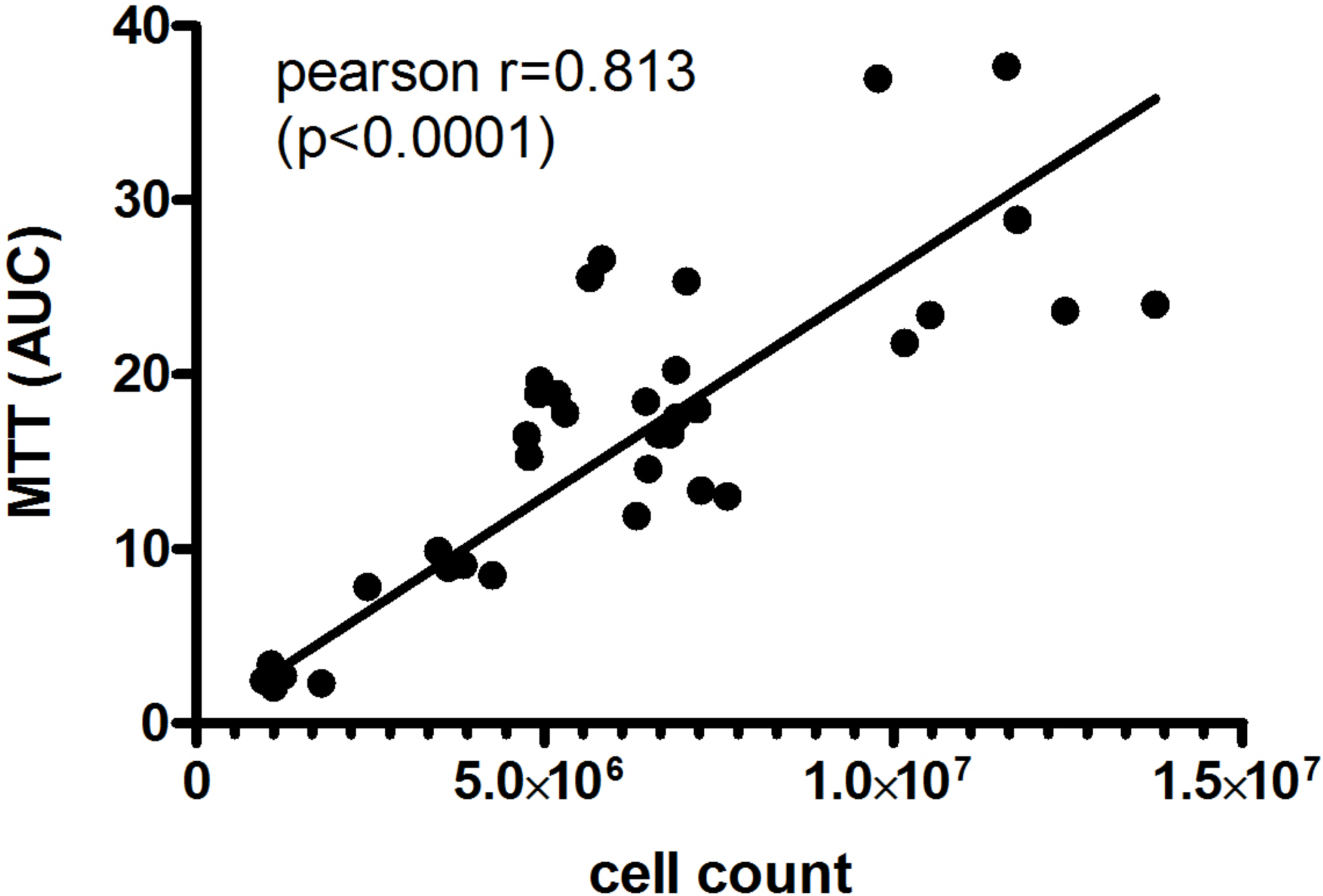
Linear relation between cell counts and MTT measurements. Cell growth was recorded by two different methods (MTT assay and cell counting) and plotted on a linear x-y graph. Both methods were significantly correlated (p<0.0001, Pearson r=0.813).

## Supplementary Table captions

**S1.**
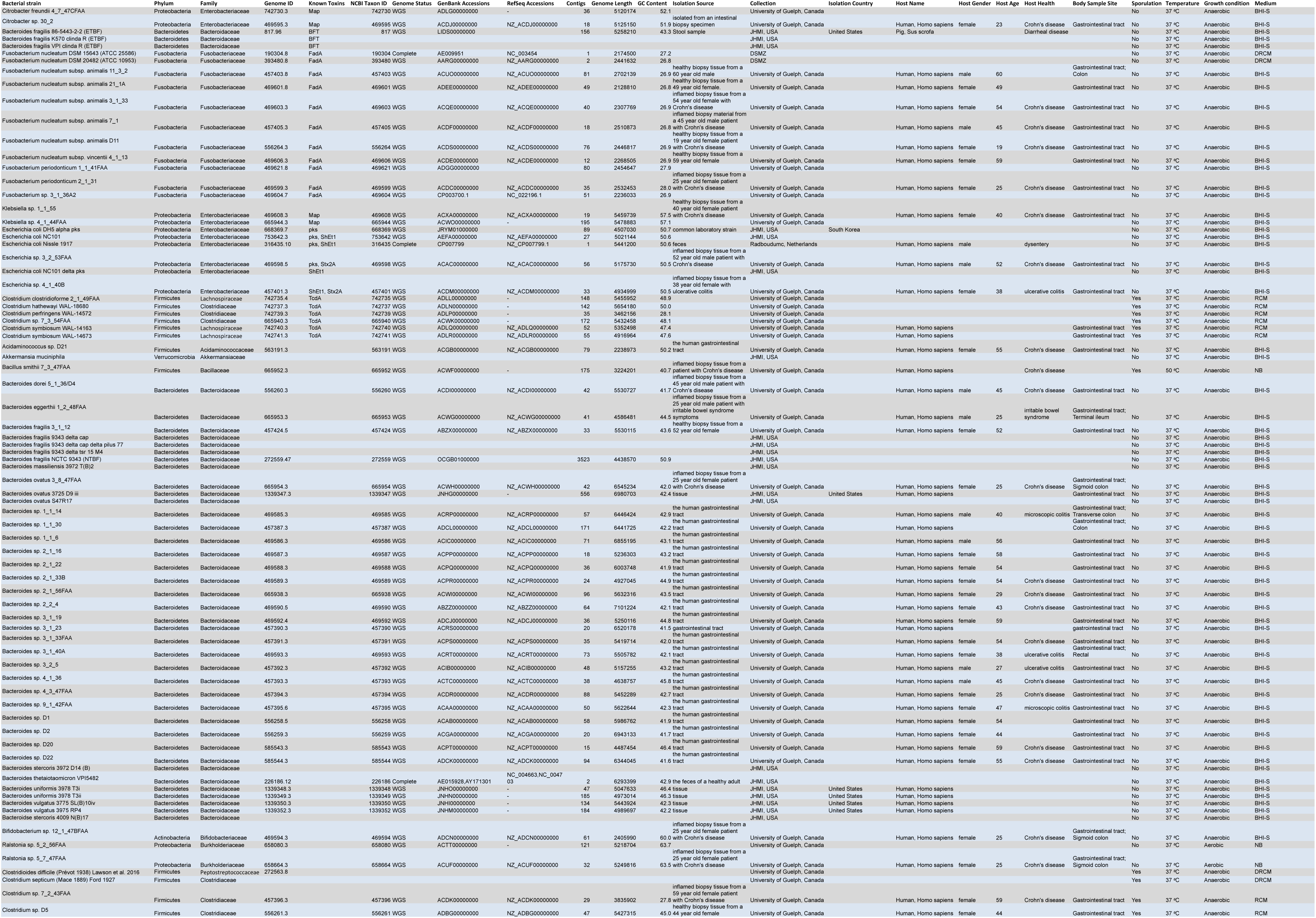
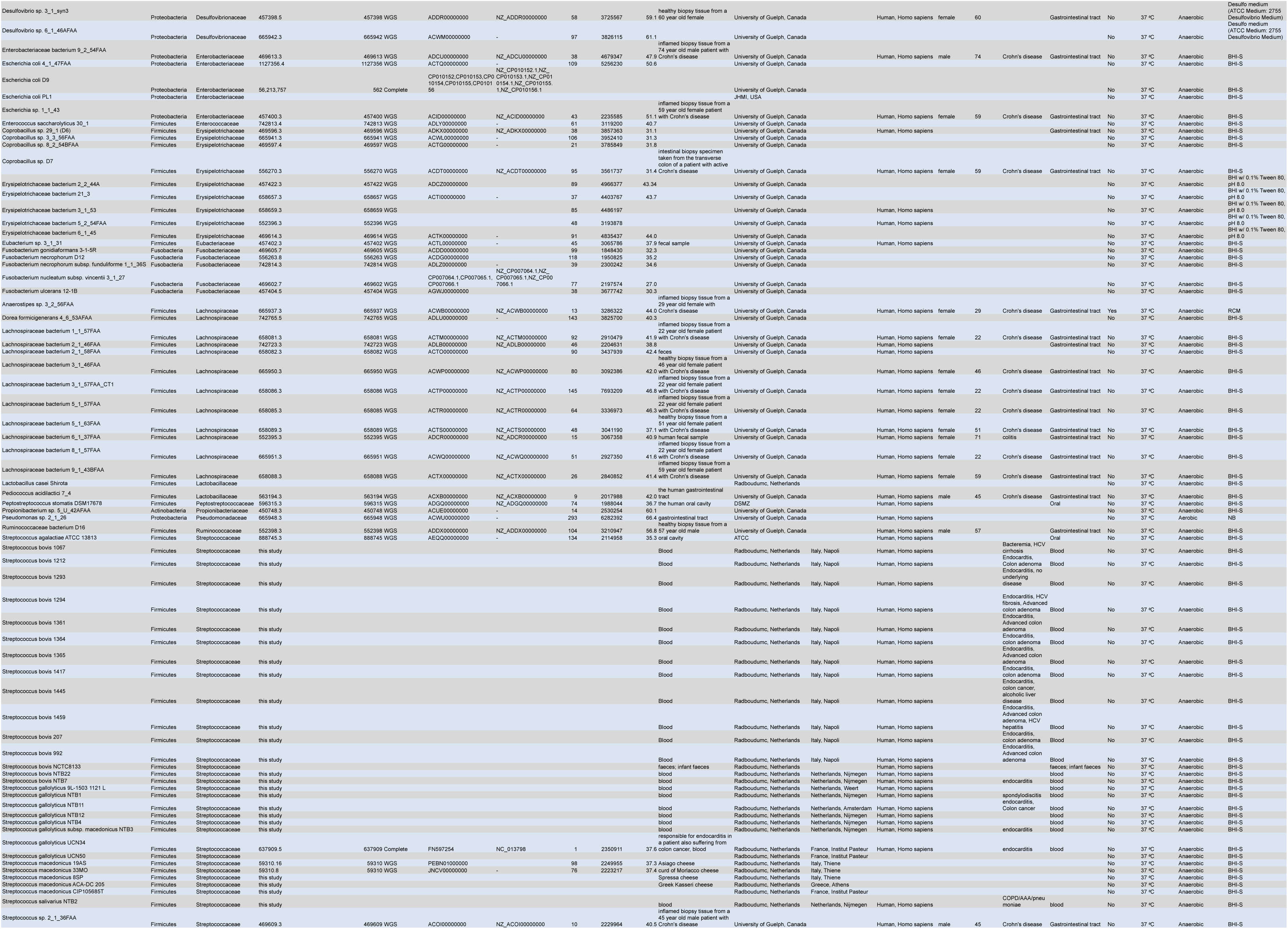
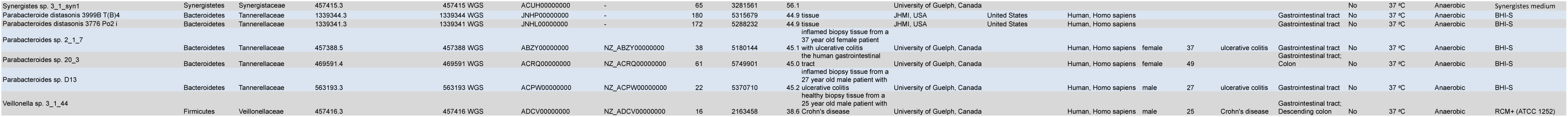
Bacterial strains used in this study.

**S2.**
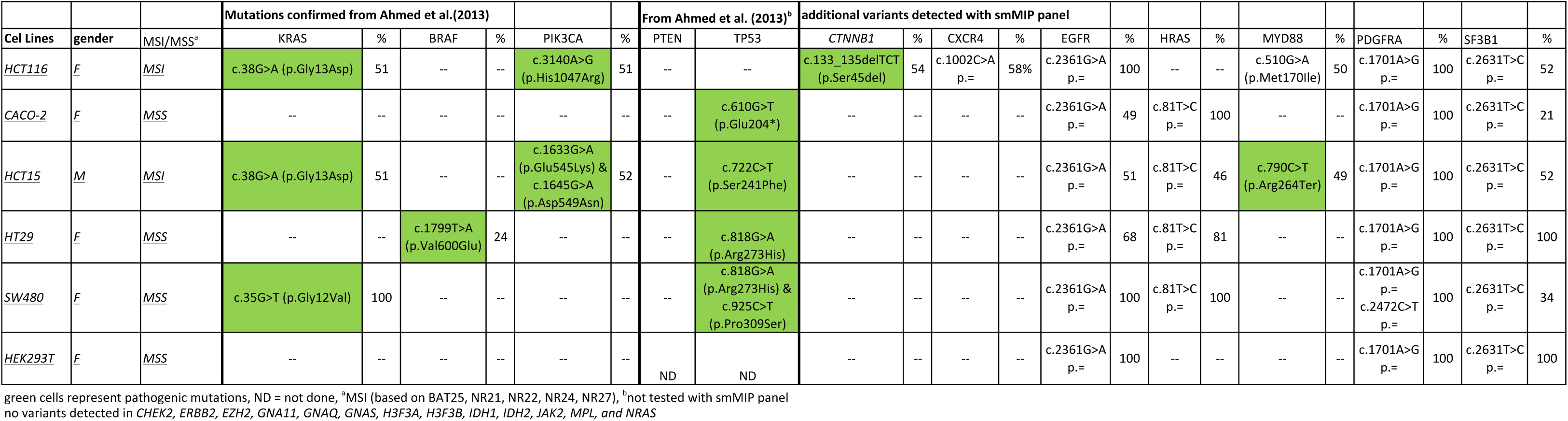
Cancer mutational profile of six human cell lines used in this study.

**S3.**
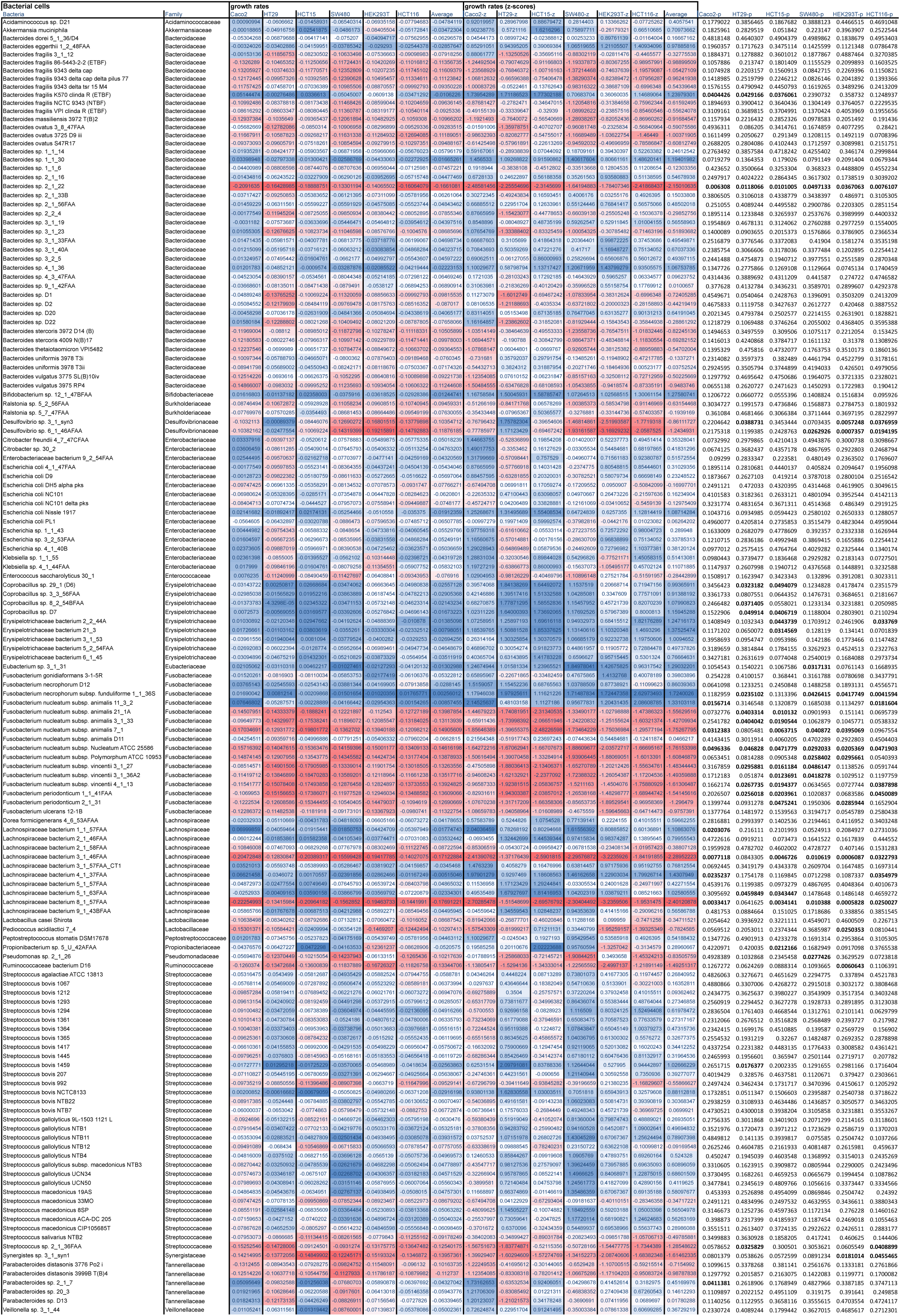
Growth rate scores, z-scores, and p-values measured from human cells incubated with bacterial cells.

**S4.**
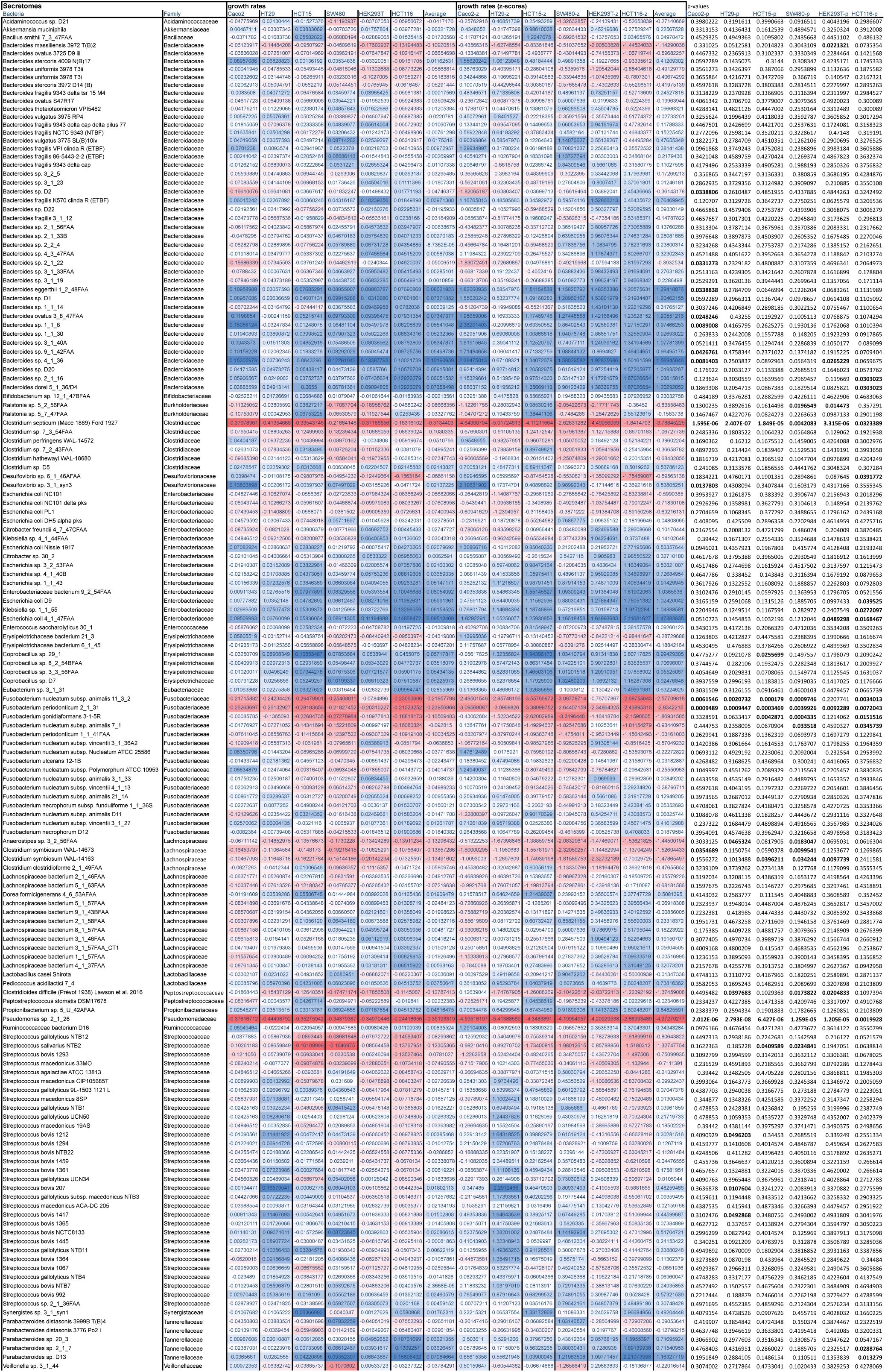
Growth rate scores, z-scores, and p-values measured from human cells incubated with bacterial secretomes.

**S5.**
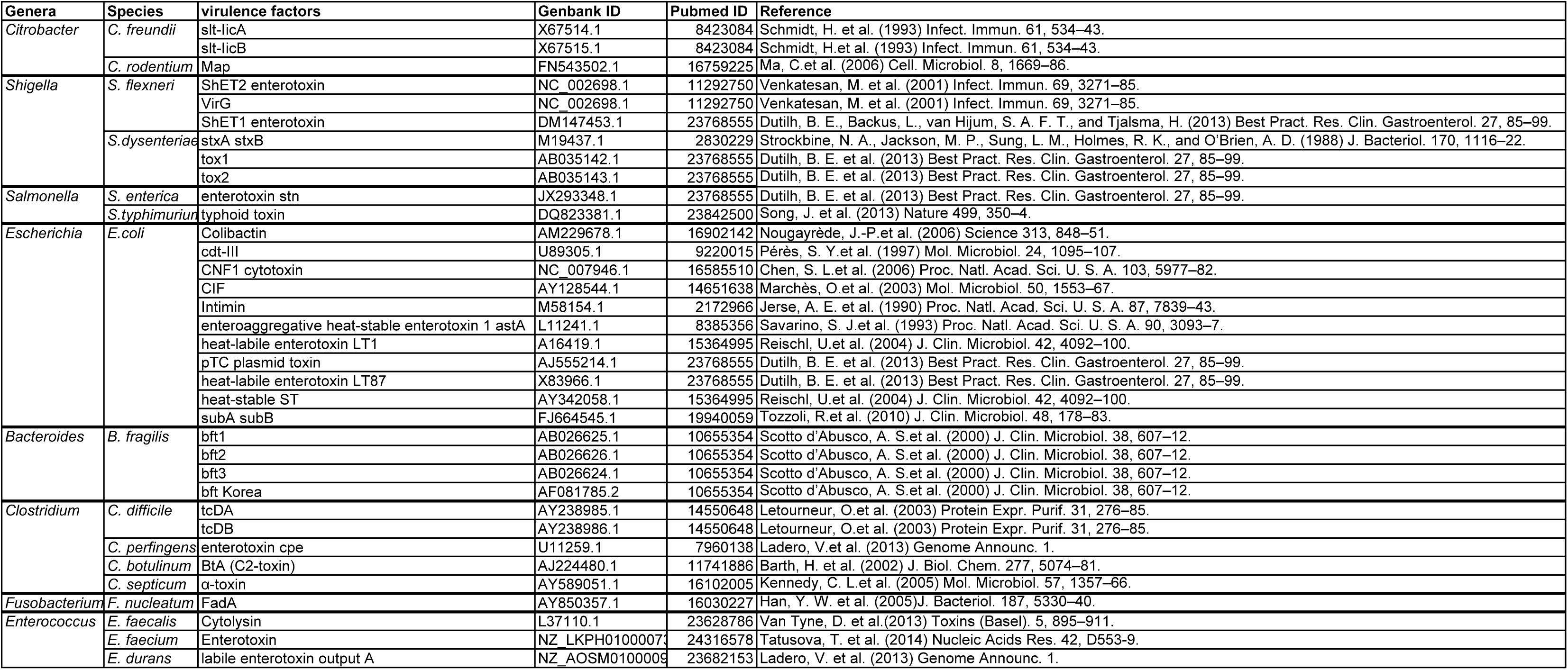
Literature summary of microbial virulence factors potentially associated to cancer.

**S6.**
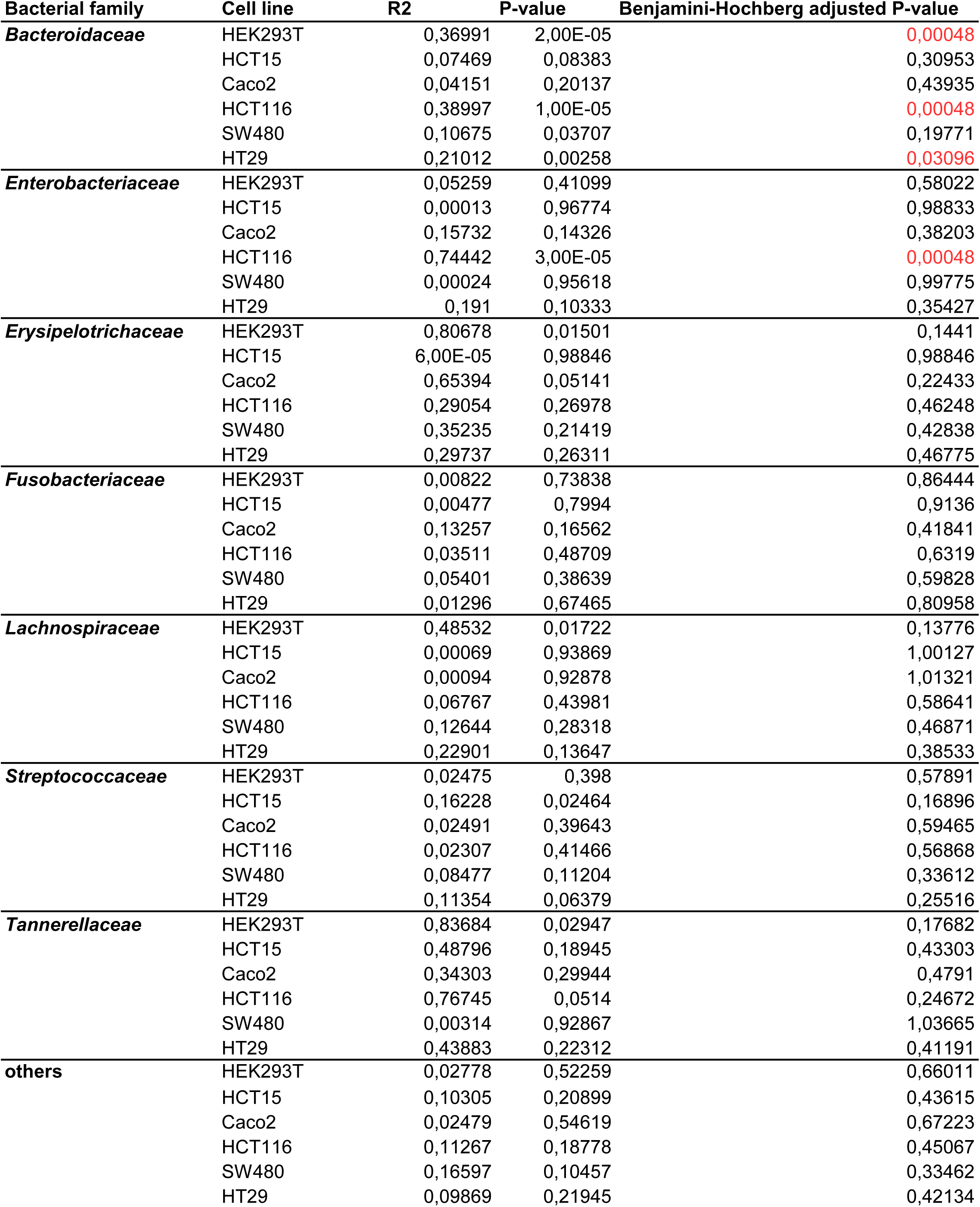
Correlation between growth rate scores of cells and secretomes.

**S7.**
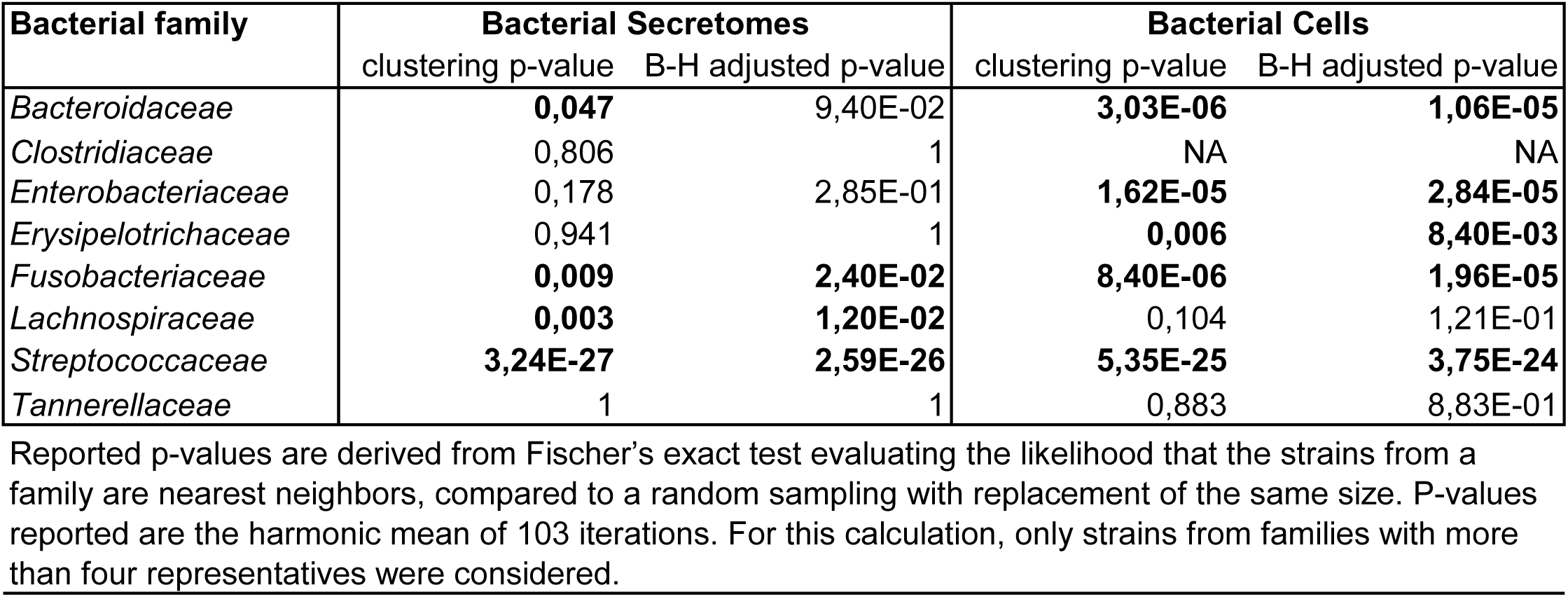
Statistical significance analysis of family-specific clustering of the growth rate scores.

**S8.**
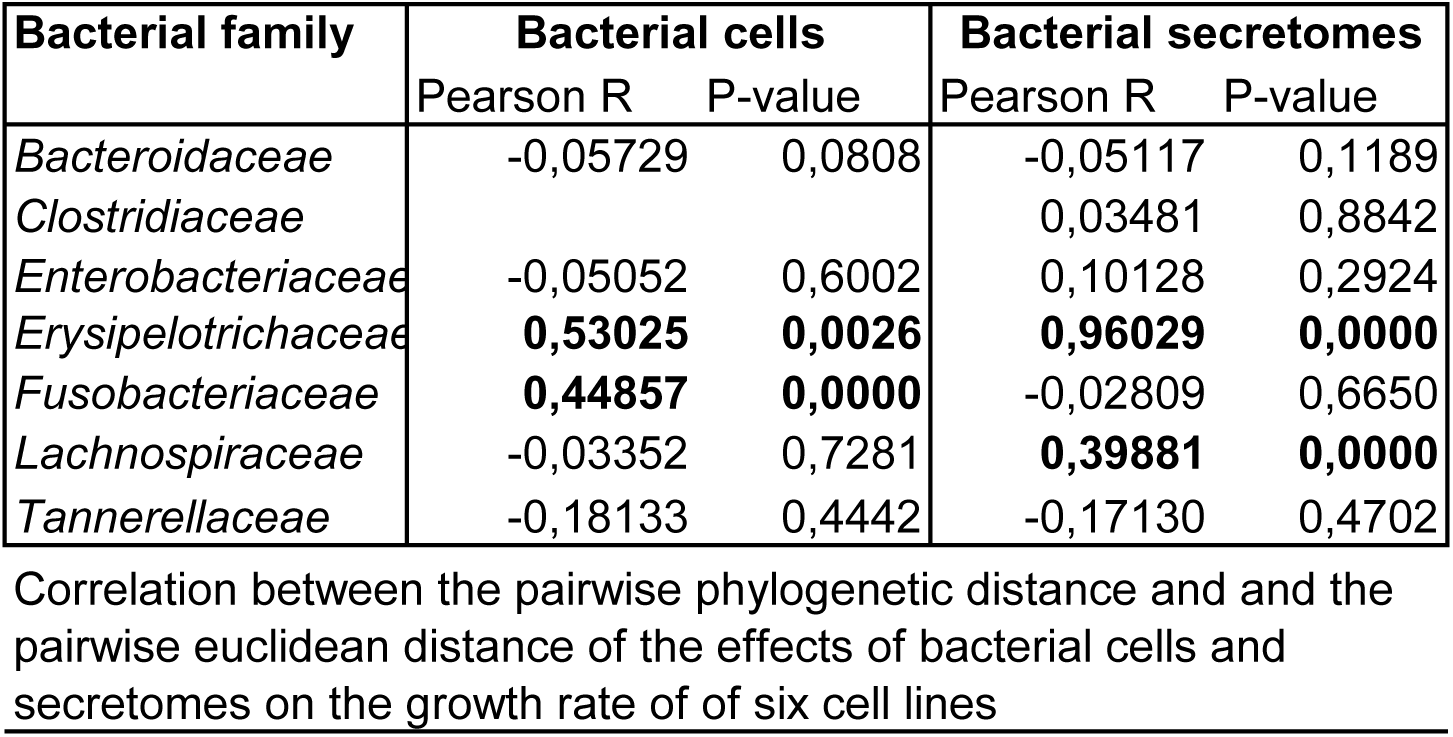
Correlation between the pairwise phylogenetic distance between bacterial strains used in this study and the pairwise Euclidean distance of the growth rate scores.

**S9.**
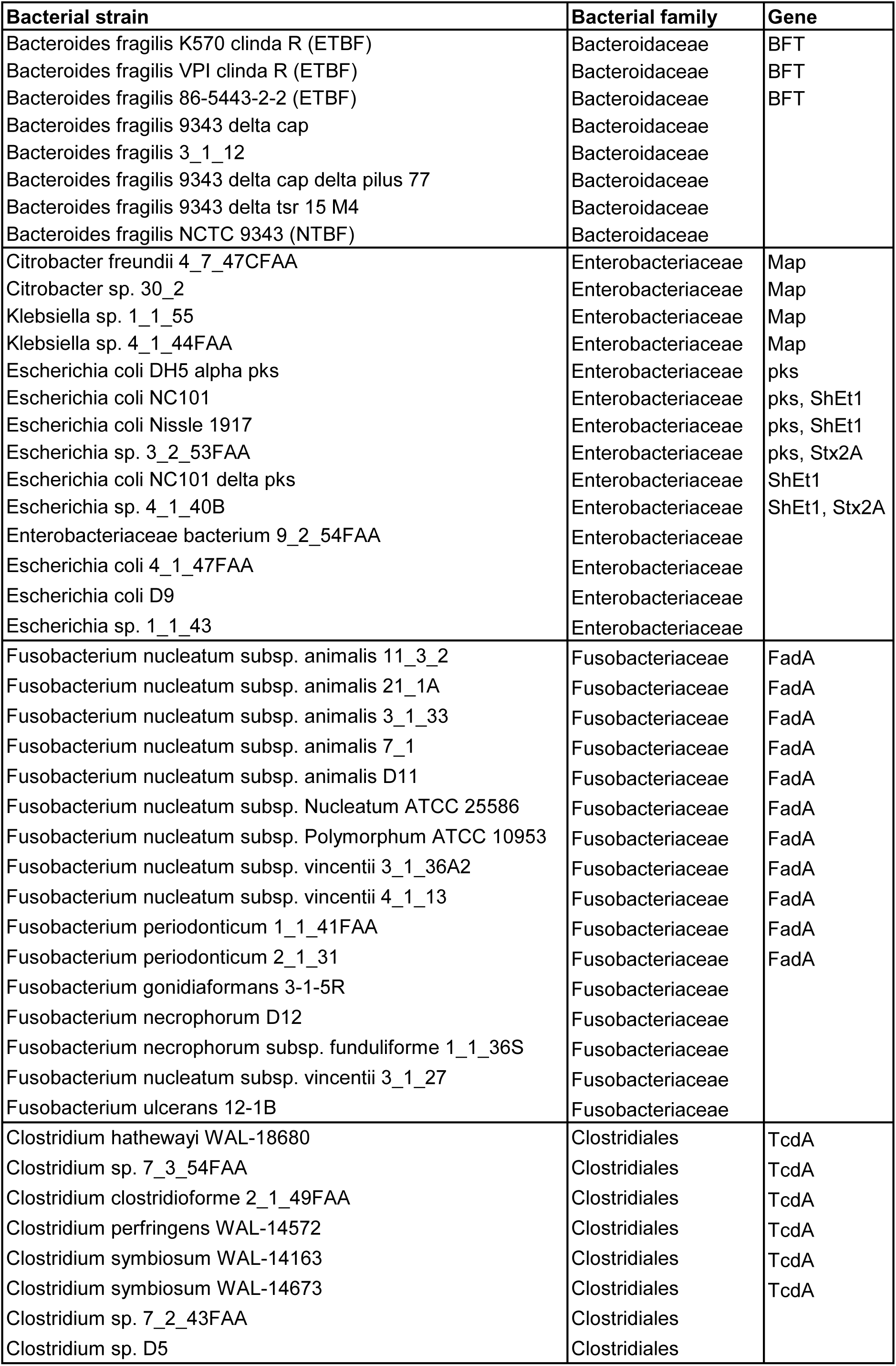
Distribution of genes coding for virulence factors in the genomes of the bacteria used in this study. The TcdA toxin was present in bacteria of the *Clostridiales* order while other toxins were present within bacterial families.

**S10.**
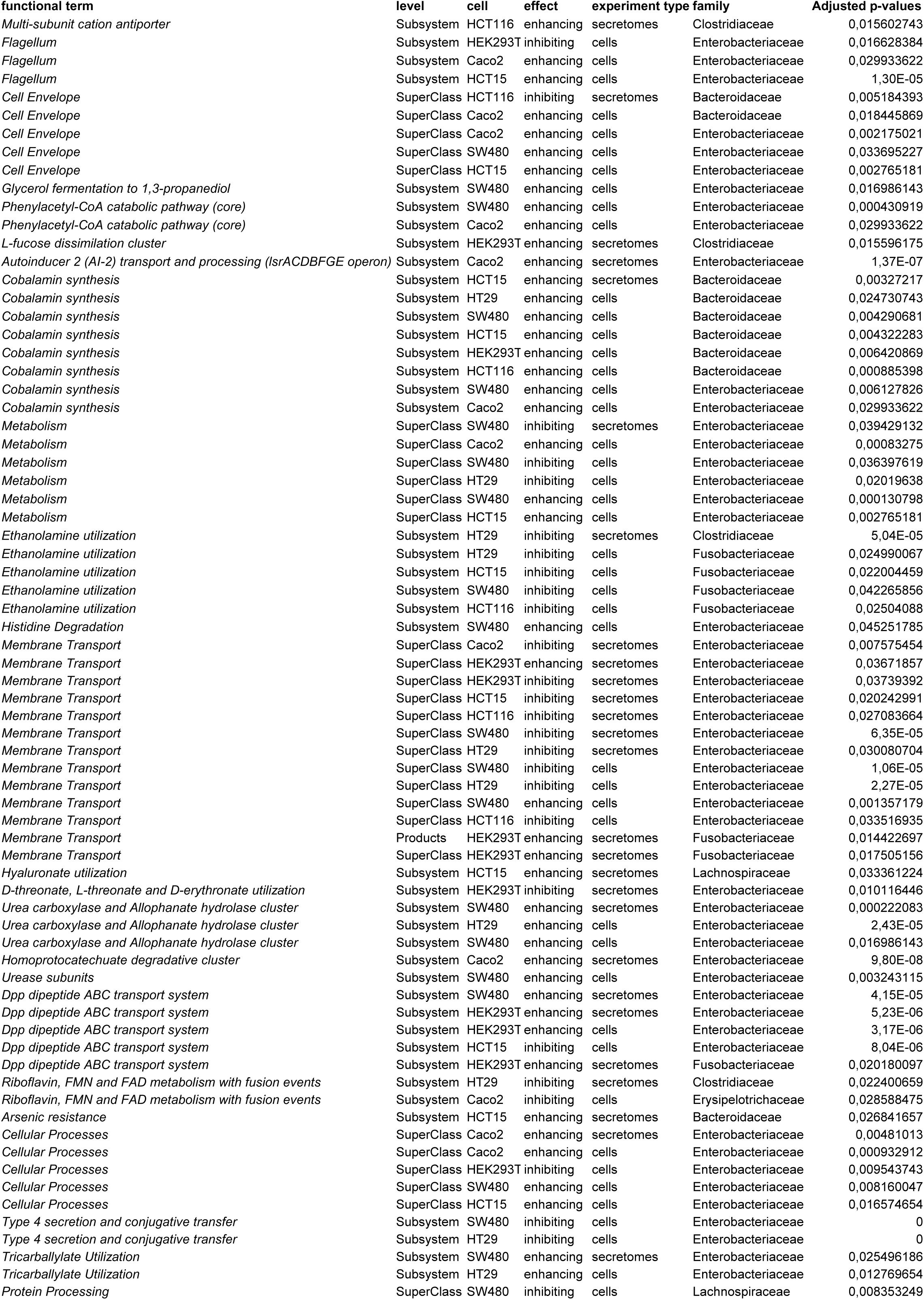
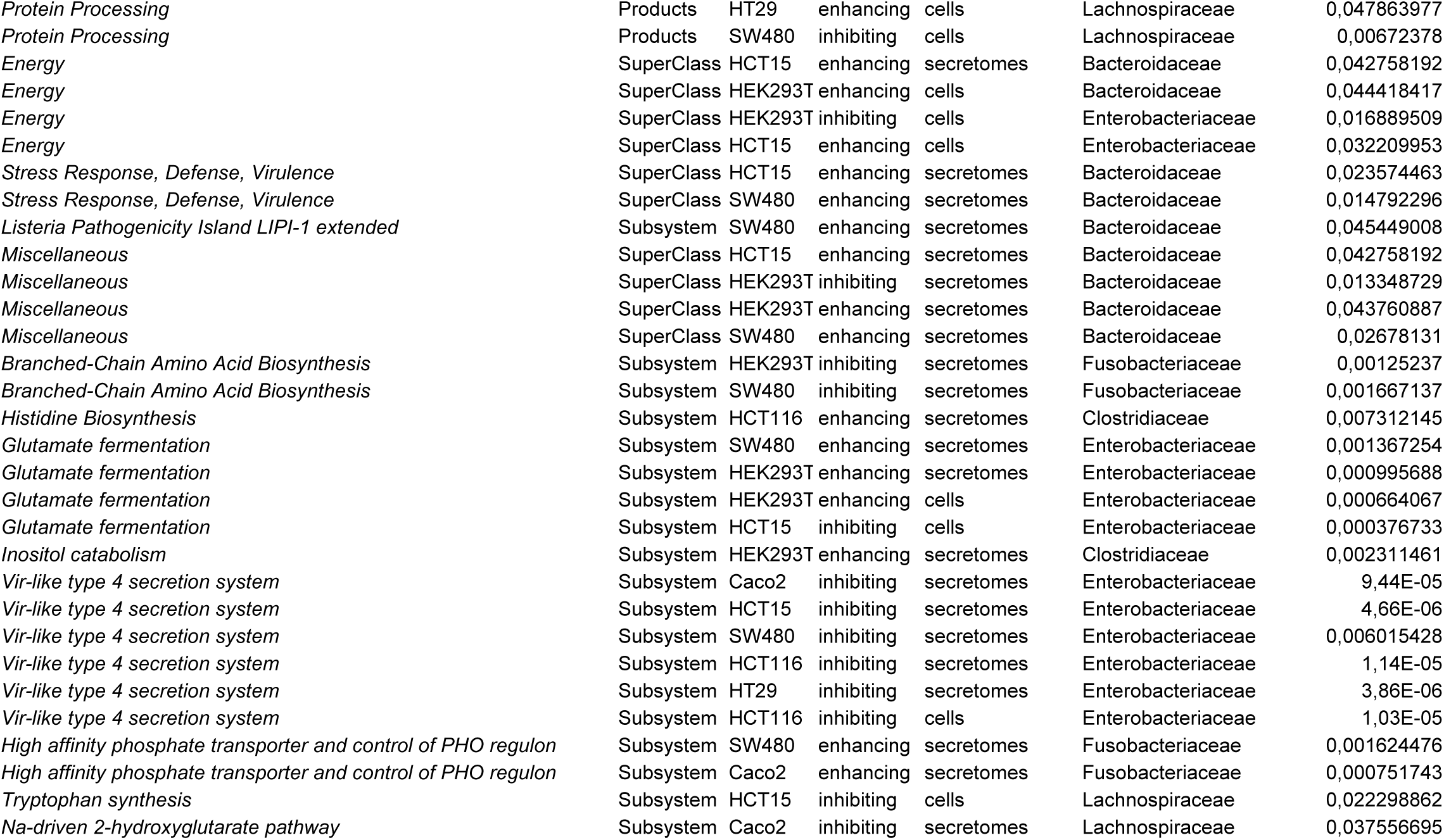
Functional genomic terms significantly associated to growth rate scores within bacterial families.

## REFERENCES

1. R. Sender, S. Fuchs, R. Milo, Revised Estimates for the Number of Human and Bacteria Cells in the Body. PLoS biology 14, e1002533–e1002533 (2016).

2. M. Rajilić-Stojanović, W. M. de Vos, The first 1000 cultured species of the human gastrointestinal microbiota. FEMS microbiology reviews 38, 996–1047 (2014).

3. J. Qin et al., A human gut microbial gene catalogue established by metagenomic sequencing. Nature 464, 59–65 (2010).

4. M. Arnold et al., Global patterns and trends in colorectal cancer incidence and mortality. Gut 66, 683–691 (2017).

5. R. Gao et al., Mucosa-associated microbiota signature in colorectal cancer. European journal of clinical microbiology & infectious diseases : official publication of the European Society of Clinical Microbiology 36, 2073–2083 (2017).

6. H. Tjalsma, A. Boleij, J. R. Marchesi, B. E. Dutilh, A bacterial driver-passenger model for colorectal cancer: beyond the usual suspects. Nature reviews. Microbiology 10, 575–582 (2012).

7. Z. Gao, B. Guo, R. Gao, Q. Zhu, H. Qin, Microbiota disbiosis is associated with colorectal cancer. Frontiers in microbiology 6, 20 (2015).

8. D. Hanahan, R. A. Weinberg, Hallmarks of cancer: the next generation. Cell 144, 646–674 (2011).

9. J. R. Marchesi et al., Towards the human colorectal cancer microbiome. PloS one 6, e20447–e20447 (2011).

10. J. Wirbel et al., Meta-analysis of fecal metagenomes reveals global microbial signatures that are specific for colorectal cancer. Nature medicine 25, 679–689 (2019).

11. K. Xu, B. Jiang, Analysis of Mucosa-Associated Microbiota in Colorectal Cancer. Medical science monitor : international medical journal of experimental and clinical research 23, 4422–4430 (2017).

12. M. S. Donia, A Toolbox for Microbiome Engineering. Cell systems 1, 21–23 (2015).

13. J. L. Foo, H. Ling, Y. S. Lee, M. W. Chang, Microbiome engineering: Current applications and its future. Biotechnology journal 12 (2017).

14. B. Garcia-Jimenez, T. de la Rosa, M. D. Wilkinson, MDPbiome: microbiome engineering through prescriptive perturbations. Bioinformatics (Oxford, England) 34, i838–i847 (2018).

15. J. L. Sonnenburg, Microbiome engineering. Nature 518, S10 (2015).

16. N. Bernardes, R. Seruca, A. M. Chakrabarty, A. M. Fialho, Microbial-based therapy of cancer: current progress and future prospects. Bioengineered bugs 1, 178–190 (2010).

17. S. Song, M. S. Vuai, M. Zhong, The role of bacteria in cancer therapy - enemies in the past, but allies at present. Infectious agents and cancer 13, 9 (2018).

18. K. Lukasiewicz, M. Fol, Microorganisms in the Treatment of Cancer: Advantages and Limitations. Journal of immunology research 2018, 2397808 (2018).

19. N. S. Forbes et al., White paper on microbial anti-cancer therapy and prevention. Journal for immunotherapy of cancer 6, 78 (2018).

20. M. G. Kramer, M. Masner, F. A. Ferreira, R. M. Hoffman, Bacterial Therapy of Cancer: Promises, Limitations, and Insights for Future Directions. Frontiers in microbiology 9, 16 (2018).

21. X. J. Shen et al., Molecular characterization of mucosal adherent bacteria and associations with colorectal adenomas. Gut microbes 1, 138–147 (2010).

22. W. Chen, F. Liu, Z. Ling, X. Tong, C. Xiang, Human intestinal lumen and mucosa-associated microbiota in patients with colorectal cancer. PloS one 7, e39743 (2012).

23. B. Flemer et al., Tumour-associated and non-tumour-associated microbiota in colorectal cancer. Gut 66, 633–643 (2017).

24. G. Zeller et al., Potential of fecal microbiota for early-stage detection of colorectal cancer. Molecular systems biology 10, 766 (2014).

25. T. Wang et al., Structural segregation of gut microbiota between colorectal cancer patients and healthy volunteers. The ISME journal 6, 320–329 (2012).

26. V. L. Hale et al., Synthesis of multi-omic data and community metabolic models reveals insights into the role of hydrogen sulfide in colon cancer. Methods (San Diego, Calif.) 149, 59–68 (2018).

27. A. Boleij et al., The Bacteroides fragilis toxin gene is prevalent in the colon mucosa of colorectal cancer patients. Clinical infectious diseases : an official publication of the Infectious Diseases Society of America 60, 208–215 (2015).

28. A. D. Kostic et al., Genomic analysis identifies association of Fusobacterium with colorectal carcinoma. Genome research 22, 292–298 (2012).

29. A. Boleij et al., Bacterial responses to a simulated colon tumor microenvironment. Molecular & cellular proteomics : MCP 11, 851–862 (2012).

30. H. Tjalsma, A. Bolhuis, J. D. Jongbloed, S. Bron, J. M. van Dijl, Signal peptide-dependent protein transport in Bacillus subtilis: a genome-based survey of the secretome. Microbiology and molecular biology reviews : MMBR 64, 515–547 (2000).

31. M. R. Rubinstein et al., Fusobacterium nucleatum promotes colorectal carcinogenesis by modulating E-cadherin/beta-catenin signaling via its FadA adhesin. Cell host & microbe 14, 195–206 (2013).

32. Y. Fardini et al., Fusobacterium nucleatum adhesin FadA binds vascular endothelial cadherin and alters endothelial integrity. Molecular microbiology 82, 1468–1480 (2011).

33. Y. Chen et al., Invasive Fusobacterium nucleatum activates beta-catenin signaling in colorectal cancer via a TLR4/P-PAK1 cascade. Oncotarget 8, 31802–31814 (2017).

34. M. S. Donnenberg et al., Role of the eaeA gene in experimental enteropathogenic Escherichia coli infection. The Journal of clinical investigation 92, 1412–1417 (1993).

35. A. E. Jerse, J. Yu, B. D. Tall, J. B. Kaper, A genetic locus of enteropathogenic Escherichia coli necessary for the production of attaching and effacing lesions on tissue culture cells. Proceedings of the National Academy of Sciences of the United States of America 87, 7839–7843 (1990).

36. O. D. Maddocks, A. J. Short, M. S. Donnenberg, S. Bader, D. J. Harrison, Attaching and effacing Escherichia coli downregulate DNA mismatch repair protein in vitro and are associated with colorectal adenocarcinomas in humans. PloS one 4, e5517 (2009).

37. K. J. Rhee et al., Induction of persistent colitis by a human commensal, enterotoxigenic Bacteroides fragilis, in wild-type C57BL/6 mice. Infection and immunity 77, 1708–1718 (2009).

38. S. Wu, P. J. Morin, D. Maouyo, C. L. Sears, Bacteroides fragilis enterotoxin induces c-Myc expression and cellular proliferation. Gastroenterology 124, 392–400 (2003).

39. G. Cuevas-Ramos et al., Escherichia coli induces DNA damage in vivo and triggers genomic instability in mammalian cells. Proceedings of the National Academy of Sciences of the United States of America 107, 11537–11542 (2010).

40. J. C. Arthur et al., Intestinal inflammation targets cancer-inducing activity of the microbiota. Science (New York, N.Y.) 338, 120–123 (2012).

41. M. R. Wilson et al., The human gut bacterial genotoxin colibactin alkylates DNA. Science (New York, N.Y.) 363 (2019).

42. A. R. Wattam et al., Improvements to PATRIC, the all-bacterial Bioinformatics Database and Analysis Resource Center. Nucleic acids research 45, D535–d542 (2017).

43. A. Boleij, M. M. van Gelder, D. W. Swinkels, H. Tjalsma, Clinical Importance of Streptococcus gallolyticus infection among colorectal cancer patients: systematic review and meta-analysis. Clinical infectious diseases : an official publication of the Infectious Diseases Society of America 53, 870–878 (2011).

44. G. K. Wentling, P. P. Metzger, E. J. Dozois, H. K. Chua, M. Krishna, Unusual bacterial infections and colorectal carcinoma--Streptococcus bovis and Clostridium septicum: report of three cases. Diseases of the colon and rectum 49, 1223–1227 (2006).

45. R. E. Schaaf et al., Clostridium septicum infection associated with colonic carcinoma and hematologic abnormality. Radiology 137, 625–627 (1980).

46. J. S. Sidhu, A. Mandal, J. Virk, V. Gayam, Early Detection of Colon Cancer Following Incidental Finding of Clostridium septicum Bacteremia. Journal of investigative medicine high impact case reports 7, 2324709619832050 (2019).

47. Y. Zheng et al., Clostridium difficile colonization in preoperative colorectal cancer patients. Oncotarget 8, 11877–11886 (2017).

48. M. H. Fukugaiti et al., High occurrence of Fusobacterium nucleatum and Clostridium difficile in the intestinal microbiota of colorectal carcinoma patients. Brazilian journal of microbiology : [publication of the Brazilian Society for Microbiology] 46, 1135–1140 (2015).

49. K. Yamazaki et al., The effect of an oral administration of Lactobacillus casei strain shirota on azoxymethane-induced colonic aberrant crypt foci and colon cancer in the rat. Oncology reports 7, 977–982 (2000).

50. A. Kato-Kataoka et al., Fermented Milk Containing Lactobacillus casei Strain Shirota Preserves the Diversity of the Gut Microbiota and Relieves Abdominal Dysfunction in Healthy Medical Students Exposed to Academic Stress. Applied and environmental microbiology 82, 3649–3658 (2016).

51. T. Morishita, T. Fukada, M. Shirota, T. Yura, Genetic basis of nutritional requirements in Lactobacillus casei. Journal of bacteriology 120, 1078–1084 (1974).

52. M. Derrien, E. E. Vaughan, C. M. Plugge, W. M. de Vos, Akkermansia muciniphila gen. nov., sp. nov., a human intestinal mucin-degrading bacterium. International journal of systematic and evolutionary microbiology 54, 1469–1476 (2004).

53. C. M. Dejea et al., Patients with familial adenomatous polyposis harbor colonic biofilms containing tumorigenic bacteria. Science (New York, N.Y.) 359, 592–597 (2018).

54. T. L. Weir et al., Stool microbiome and metabolome differences between colorectal cancer patients and healthy adults. PloS one 8, e70803 (2013).

55. C. Dingemanse et al., Akkermansia muciniphila and Helicobacter typhlonius modulate intestinal tumor development in mice. Carcinogenesis 36, 1388–1396 (2015).

56. A. Boleij et al., Novel clues on the specific association of Streptococcus gallolyticus subsp gallolyticus with colorectal cancer. The Journal of infectious diseases 203, 1101–1109 (2011).

57. M. F. Tripodi et al., Streptococcus bovis endocarditis and its association with chronic liver disease: an underestimated risk factor. Clinical infectious diseases : an official publication of the Infectious Diseases Society of America 38, 1394–1400 (2004).

58. M. F. Tripodi, R. Fortunato, R. Utili, M. Triassi, R. Zarrilli, Molecular epidemiology of Streptococcus bovis causing endocarditis and bacteraemia in Italian patients. Clinical microbiology and infection : the official publication of the European Society of Clinical Microbiology and Infectious Diseases 11, 814–819 (2005).

59. Y. Ogawa et al., The genome of Erysipelothrix rhusiopathiae, the causative agent of swine erysipelas, reveals new insights into the evolution of firmicutes and the organism’s intracellular adaptations. Journal of bacteriology 193, 2959–2971 (2011).

60. R. Taddese et al., Bacterial Zombies and Ghosts: Production of Inactivated Gram-Positive and Gram-Negative Species with Preserved Cellular Morphology and Cytoplasmic Content. bioRxiv 10.1101/458158, 458158 (2018).

61. D. Ahmed et al., Epigenetic and genetic features of 24 colon cancer cell lines. Oncogenesis 2, e71 (2013).

62. A. Eijkelenboom et al., Reliable Next-Generation Sequencing of Formalin-Fixed, Paraffin-Embedded Tissue Using Single Molecule Tags. The Journal of molecular diagnostics : JMD 18, 851–863 (2016).

63. T. Mosmann, Rapid colorimetric assay for cellular growth and survival: application to proliferation and cytotoxicity assays. Journal of immunological methods 65, 55–63 (1983).

64. L. H. van der Maaten, Geoffrey, Visualizing Data using t-SNE. Journal of Machine Learning Research 9, 2579--2605 (2008).

65. J. L. Bentley, Multidimensional binary search trees used for associative searching. Commun. ACM 18, 509–517 (1975).

66. R. K. Aziz et al., The RAST Server: rapid annotations using subsystems technology. BMC genomics 9, 75 (2008).

67. Anonymous, “Poisson Distribution” in Univariate Discrete Distributions. 10.1002/0471715816.ch4, pp. 156–207.

68. Y. Xi, Z. Ma, H. Zhang, M. Yuan, L. Wang, Effects of Clostridium difficile toxin A on K562/A02 cell proliferation, apoptosis and multi-drug resistance. Oncology letters 15, 4215–4220 (2018).

69. G. Dalmasso, A. Cougnoux, J. Delmas, A. Darfeuille-Michaud, R. Bonnet, The bacterial genotoxin colibactin promotes colon tumor growth by modifying the tumor microenvironment. Gut Microbes 5, 675–680 (2014).

70. T. Fais, J. Delmas, N. Barnich, R. Bonnet, G. Dalmasso, Colibactin: More Than a New Bacterial Toxin. Toxins 10 (2018).

71. P. Dean, B. Kenny, The effector repertoire of enteropathogenic E. coli: ganging up on the host cell. Current opinion in microbiology 12, 101–109 (2009).

72. A. P. Singh et al., Enteropathogenic E. coli effectors EspF and Map independently disrupt tight junctions through distinct mechanisms involving transcriptional and post-transcriptional regulation. Scientific reports 8, 3719 (2018).

73. T. M. Wassenaar, E. coli and colorectal cancer: a complex relationship that deserves a critical mindset. Critical reviews in microbiology 44, 619–632 (2018).

74. N. Bossuet-Greif et al., The Colibactin Genotoxin Generates DNA Interstrand Cross-Links in Infected Cells. mBio 9 (2018).

75. J. Ahn et al., Human gut microbiome and risk for colorectal cancer. Journal of the National Cancer Institute 105, 1907–1911 (2013).

76. Y. Yang et al., Fusobacterium nucleatum Increases Proliferation of Colorectal Cancer Cells and Tumor Development in Mice by Activating Toll-Like Receptor 4 Signaling to Nuclear Factor-kappaB, and Up-regulating Expression of MicroRNA-21. Gastroenterology 152, 851–866.e824 (2017).

77. M. Castellarin et al., Fusobacterium nucleatum infection is prevalent in human colorectal carcinoma. Genome research 22, 299–306 (2012).

78. C. T. Ma, H. S. Luo, F. Gao, Q. C. Tang, W. Chen, Fusobacterium nucleatum promotes the progression of colorectal cancer by interacting with E-cadherin. Oncology letters 16, 2606–2612 (2018).

79. J. Liu et al., Proteomic characterization of outer membrane vesicles from gut mucosa-derived fusobacterium nucleatum. Journal of proteomics 195, 125–137 (2019).

80. Y. W. Han et al., Identification and Characterization of a Novel Adhesin Unique to Oral Fusobacteria. Journal of bacteriology 187, 5330–5340 (2005).

81. C. L. Sears, The toxins of Bacteroides fragilis. Toxicon 39, 1737–1746 (2001).

82. C. Jans, A. Boleij, The Road to Infection: Host-Microbe Interactions Defining the Pathogenicity of Streptococcus bovis/Streptococcus equinus Complex Members. Frontiers in microbiology 9, 603 (2018).

83. A. Boleij, H. Tjalsma, The itinerary of Streptococcus gallolyticus infection in patients with colonic malignant disease. Lancet Infect Dis 13, 719–724 (2013).

84. R. Kumar et al., Streptococcus gallolyticus subsp. gallolyticus promotes colorectal tumor development. PLoS Pathog 13, e1006440 (2017).

85. J. Yang et al., Adenomatous polyposis coli (APC) differentially regulates beta-catenin phosphorylation and ubiquitination in colon cancer cells. J Biol Chem 281, 17751–17757 (2006).

86. J. G. Tate et al., COSMIC: the Catalogue Of Somatic Mutations In Cancer. Nucleic acids research 47, D941–d947 (2019).

87. G. G. Sgro et al., Bacteria-Killing Type IV Secretion Systems. Frontiers in microbiology 10, 1078 (2019).

88. B. Kiersztyn, W. Siuda, R. J. Chrost, Persistence of bacterial proteolytic enzymes in lake ecosystems. FEMS Microbiol Ecol 80, 124–134 (2012).

89. S. F. Battaglia-Hsu et al., Vitamin B12 deficiency reduces proliferation and promotes differentiation of neuroblastoma cells and up-regulates PP2A, proNGF, and TACE. Proceedings of the National Academy of Sciences of the United States of America 106, 21930–21935 (2009).

90. A. G. Wexler et al., Human gut Bacteroides capture vitamin B12 via cell surface-exposed lipoproteins. Elife 7 (2018).

91. D. Patel, S. N. Witt, Ethanolamine and Phosphatidylethanolamine: Partners in Health and Disease. Oxid Med Cell Longev 2017, 4829180 (2017).

92. O. Tsoy, D. Ravcheev, A. Mushegian, Comparative genomics of ethanolamine utilization. Journal of bacteriology 191, 7157–7164 (2009).

93. K. G. Kaval, D. A. Garsin, Ethanolamine Utilization in Bacteria. mBio 9 (2018).

94. M. M. Kendall, C. C. Gruber, C. T. Parker, V. Sperandio, Ethanolamine controls expression of genes encoding components involved in interkingdom signaling and virulence in enterohemorrhagic Escherichia coli O157:H7. mBio 3 (2012).

95. M. van de Wetering et al., Prospective derivation of a living organoid biobank of colorectal cancer patients. Cell 161, 933–945 (2015).

